# Paternally expressed imprinted *Snord116* and *Peg3* regulate hypothalamic orexin neurons

**DOI:** 10.1101/820738

**Authors:** Pace Marta, Falappa Matteo, Freschi Andrea, Balzani Edoardo, Berteotti Chiara, Lo Martire Viviana, Fatemeh Kaveh, Eivind Hovig, Zoccoli Giovanna, Cerri Matteo, Amici Roberto, Urbanucci Alfonso, Tucci Valter

## Abstract

Imprinted genes are highly expressed in the hypothalamus; however, whether specific imprinted genes affect hypothalamic neuromodulators and their functions is unknown. It has been suggested that Prader-Willi syndrome (PWS), a neurodevelopmental disorder caused by lack of paternal expression at chromosome 15q11-q13, is characterised by hypothalamic insufficiency. Here, we investigate the role of the paternally expressed *Snord116* gene within the context of sleep and metabolic abnormalities of PWS, and we report a novel role of this imprinted gene in the function and organisation of the two main neuromodulatory systems of the lateral hypothalamus (LH), namely, the orexin (OX) and melanin concentrating hormone (MCH) systems. We observe that the dynamics between neuronal discharge in the LH and the sleep-wake states of mice with paternal deletion of *Snord116* (PWScr^m+/p−^) are compromised. This abnormal state-dependent neuronal activity is paralleled by a significant reduction in OX neurons in the LH of mutants. Therefore, we propose that an imbalance between OX- and MCH-expressing neurons in the LH of mutants reflects a series of deficits manifested in the PWS, such as dysregulation of rapid eye movement (REM) sleep, food intake and temperature control.

**Highlights:**

- *Snord116* regulates neuronal activity in the lateral hypothalamus (LH), which is time-locked with cortical states of sleep.
- Loss of *Snord116* reduces orexin neurons in the LH and affects sleep homeostasis and thermoregulation in mice.
- *Snord116* and *Peg3* independently control orexin expression in the LH.
- Paternally expressed alleles maximize the patrilineal effects in the control of REM sleep by the LH in mammals.

## Introduction

Both maternally and paternally derived genes are essential for survival beyond post-fertilization; these genes differentially affect embryonic brain development and, consequently, postnatal and adult physiology. In particular, paternally derived genes are thought to control the organization of the subcortical limbic system [1]. For example, androgenetic (two paternal copies) cells are mainly distributed in the hypothalamus, although the specific impact of such parental genetic information on hypothalamic functions remains unknown.

The hypothalamus is an ancient structure that orchestrates primitive physiological processes for survival [2], such as motivated behaviours for feeding and drinking, the regulation of body temperature and the switch between sleep and wakefulness. Therefore, a number of paternally expressed genes that are highly expressed in the hypothalamus are potential regulators of mammalian sleep and sleep-mediated metabolism [3]. To this end, over the last decade, we have demonstrated that parent-of-origin imprinted genes exert a pivotal role in the control of sleep physiology and feeding behaviour [3, 4]

Among the pathological conditions that depend on genomic imprinting defects, Prader–Willi syndrome (PWS) is the neurodevelopmental disorder that best describes the link among sleep, metabolism and imprinted genes. PWS results from the loss of a cluster of paternally expressed genes on the chromosome 15q11-q13 region, many of which are highly expressed in the hypothalamus and are characterized by sleep-wake (e.g., rapid-eye-movement, REM, alterations) and metabolic (e.g., hyperphagia) abnormalities. All these symptoms are generally associated with hypothalamic insufficiency [5–7].

We have previously described that microdeletion of the small nuclear ribonucleic acid (RNA)-116 (*SNORD116*) cluster within the PWS locus induces REM and temperature dysregulations in mice and human subjects [5]. Specifically, the deletion of *Snord116* in mice causes an EEG profile characterized by the intrusion of REM sleep episodes into the transition between wakefulness and sleep accompanied by an increase in body temperature. REM sleep intrusions have been reported in several clinical studies in which PWS subjects manifest narcolepsy and express symptoms such as sleep attacks during active wakefulness, cataplexy (a transient loss of muscle tone during wakefulness), sleep paralysis and sleep fragmentation [8]. Narcolepsy is a sleep condition that causes the loss of hypothalamic OX neurons (also known as hypocretin; HCRT)[9], and previous studies have observed that subjects with PWS show OX alterations [10–12].

OX neurons are located in the lateral hypothalamus (LH), where this class of neurons promotes wakefulness [13] by facilitating the release of other arousal-promoting brain neuromodulators (i.e., noradrenaline, histamine and acetylcholine) [14]. However, in the LH, OX neurons are intermingled with a group of neurons that release the melanin-concentrating hormone (MCH) and promote sleep, and these neurons are active mainly during REM sleep [15]. Both OX and MCH neurons project widely throughout the brain, exerting antagonistic actions on brain states and energy balance. However, whether these two groups of neurons of the LH exert abnormal control over sleep-wake cycles, feeding and temperature in the PWS remains unclear.

In this study, we tested the hypothesis that the paternally expressed *Snord116* modulates the neuromodulatory systems of LH, therefore controlling sleep, feeding and temperature. We found that mice with paternal deletion of *Snord116* have altered dynamics in how neuronal activity of the LH is associated with cortical states. We report that in mutant mice, compared with wild-type mice, a high proportion of LH neurons do not uniquely respond to cortical states, such as those occurring in sleep, wakefulness or feeding. This altered modulation between cortical states and subcortical neuronal activity in *Snord116*-deleted mice is compatible with a loss of OX-expressing neurons in the LH, while MCH-expressing neurons remained unaffected, thereby creating an imbalance between the two systems. We also report, for the first time, a link between *Snord116* and a different paternally imprinted gene, *Peg3*, which plays a pivotal role in the control of the hypothalamic OX neuromodulatory system.

## Results

### Loss of paternal *Snord116* alters neuronal dynamics in the LH associated with sleep homeostasis

To investigate whether the firing pattern of neurons within the LH manifests signs of paternal-dependent subcortical regulation throughout different arousal states of the brain, we studied mice with paternal deletion of the *Snord116* gene [16], PWScr^m+/p−^ mice, and their wild-type littermate controls, PWScr^m+/p+^ mice. We identified neurons in the LH that are time-locked (see Methods and Figure 1A-F for full details on the procedure) to sleep (S-max) and to wake (W-max) states and to food intake (Type I, II and III). Strikingly, we observed a significant increase in S-max and a reduction in W-max neurons in PWScr^m+/p−^ mutant mice compared with controls at baseline (Figure 1G and Table S2). A more detailed analysis of B1 and B2 confirmed this distribution of the classes of LH neurons according to EEG states (Figure S1B and Table S2). Moreover, when the pressure of sleep is low (*i.e*., B1), mutant mice exhibit more REM sleep than their littermate controls (Figure S1A). This finding suggests that the pressure/need of sleep may be either a permissive or a blocking mechanism with respect to excess REM sleep due to the lack of *Snord116*. This phenomenon is confirmed by the profile of the power densities in REM sleep (Figure S1C). Indeed, in PWScr^m+/p−^ mutant mice, the theta (4.5-9 Hz) band, which is characteristic of REM sleep, is higher in both B1 and B2, but at B2, the EEG frequency phenotype is not accompanied by longer REM sleep duration, implying that the overall higher sleep pressure masks differences in microstructural aspects of electrophysiological sleep. Moreover, the delta (0.5-4.5 Hz) band presents an opposite trend in the two groups of animals; counterintuitively, reduced power is observed in mutant mice as the pressure of sleep increases (Figure S1C). Again, this phenomenon suggests abnormal homeostatic control of sleep in mutant mice.

**Figure 1.**
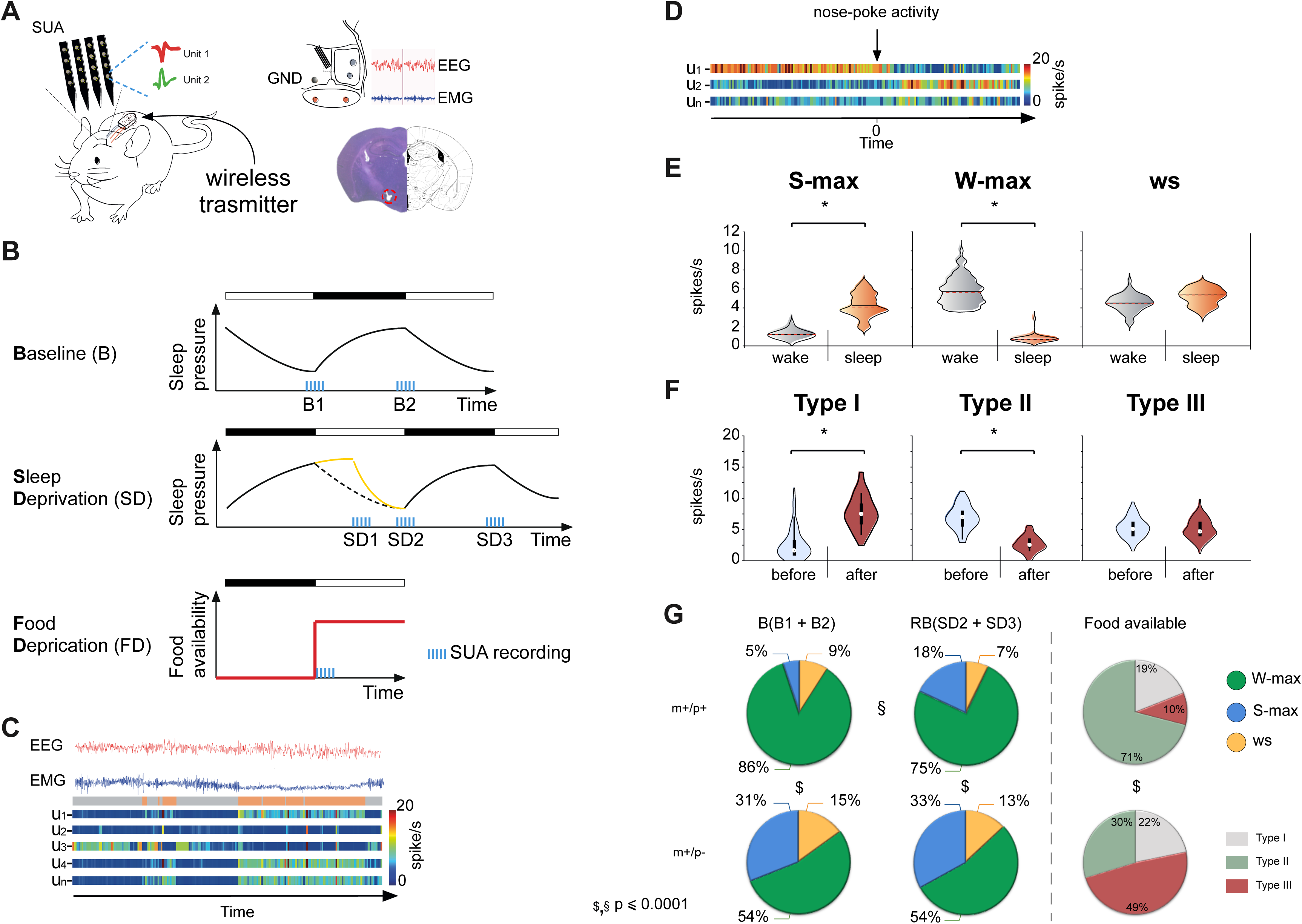
Loss of paternal *Snord116* alters neuronal dynamics in the LH associated with sleep and food. **A)** The cartoon on the left shows mice chronically implanted with a microwire array of 16 channels with an EEG-EMG wireless transmitter to record the sleep-wake cycle. On the right, a schematic representation of the mouse skull depicting EEG, EMG (red circle) and single unit activity (SUA) in the contralateral hemisphere is shown. The correct placement of the SUA electrode was histologically verified by 40-μm Nissl-stained coronal brain sections (bregma –1.10/–1.90). **B)** The panel shows a schematic representation of the experimental design used to record SUA and the sleep-wake cycle in relation to sleep pressure depicted by EEG/EMG. Recordings were performed at two different baseline (BL) time points (B1 and B2) over the first hour of the rebound period (SD1) following 6-h of total sleep deprivation (SD) and at other two time points over the 18-h recovery period (SD2 and SD3). Each recording is 2-h, except for SD1 (1-Hr). Each recording is represented by cyan bars. After 5 days of recovery from the previous SD, the animals were fasted for 12-h during the dark period and fed at the beginning of the light period. Food pellets were provided only after a nose-poke activity. SUA was continuously recorded for 2-h after being fed. The black bar indicates the dark period of the light/dark cycle, while the white bar indicates the light period of the cycle. **C)** The panel shows an example of EEG/EMG traces with the sleep stages identified (wakefulness is represented in grey, while sleep, including both NREM and REM sleep stages, is shown in orange) aligned with the firing rate recorded in the LH. The heatmaps at the bottom show the response firing rate in spikes/second from 0 Hz (blue) to 20 Hz (red). **D)** The heatmaps show the response firing rate used to classify neurons before and after food consumption (firing rate in spikes/second from 0 Hz [blue] to 20 Hz [red]). **E)** Violin plots of classified units according to the sleep-wake states in which they maximally fired according to ANOVA followed by post hoc Bonferroni correction (p < .05) see Methods. **F)** Violin plots of classified units according to their discharge related to food consumption (paired Student’s t-test of the firing rate between before and after the pellet was released, binned at 50 ms, p < 0.05). See Methods. **G)** The pie chart represents the distribution of recorded neurons according sleep-wake stage: wake (W-max), sleep (both NREM and REM sleep, S-max) and not responding (ws). The rows show the two genotypes (PWScr^m+/p+^ on the top and PWScr^m+/p−^ on the bottom), while the columns show the different recording time points, B, RB and food. According to the neuronal classification (see Methods), W-max neurons are shown in green, S-max neurons are shown in blue, and ws neurons are shown in yellow. Type I, neurons that fire maximally immediately after nose-poke activity and during feeding shown in grey; Type II, neurons that show a sharp increase in firing rate at the beginning of feeding shown in green; Type III, neurons that do not respond to food shown in red. Differences between the two genotypes are indicated by $, while differences within groups across time points are indicated by §. Significance was computed with the chi-square test; for details on the statistical analysis, see Table S2, S3 and S4. The two genotypes investigated were PWScr^m+/p−^ mice (n=4) and PWScr^m+/p+^ mice (n= 4).

Thus, we sought to explore the physiological responses of mice following 6-h of total sleep deprivation (SD) by investigating the distributions of the neuronal classes at different time points during 18 h of recovery (i.e., SD1, SD2 and SD3, Figure 1B). We observed that SD induced an increase in S-max neurons in wild-type PWScr^m+/p+^ mice, while in mutant PWScr^m+/p−^ mice, the distribution of classes of neurons remained unaltered compared with baseline (Figure 1G). This result indicates that control mice homeostatically respond to SD by modulating LH neuronal dynamics, as previously reported in the literature [17], while mutant mice lack this hypothalamic modulatory process.

Across different conditions, neurons may continue responding to the same state, may respond to different state, or may become aspecific in their activity. Within the rearrangements of neuronal modulation across different phases, an interesting observation in our study is represented by the behaviour of the non-responding neurons following SD (Figure S1A, Table S3). In wild-type mice after SD, the number of neurons of this latter class increases three times compared with the previous baseline, a phenomenon that is gradually recovered over the following 18-h after deprivation. This change comes with a cost for wake-dependent neurons in wild-type animals. In PWScr^m+/p−^ mutants, this SD effect is missing, reinforcing the observation of a lack of neuromodulation caused by the paternally derived genetic defect.

Finally, the dramatic drop in delta and theta power densities in PWScr^m+/p−^ mutant mice compared with controls at SD2 (Figure S1C) coincides with the permissive role of sleep pressure at this time of the day for the sleep defects in mutants.

### OX putative neurons are reduced in PWScr^m+/p−^ mutants compared with controls

The LH is made by a heterogeneous group of cells that express various neuropeptides, such as OX and MCH neurons. To gain insight about the heterogeneous nature of our recording units, we plotted the mean firing rate versus the mean logarithm of the EMG signals across three groups of putative neurons (W-max; S-max and ws neurons) of the whole sleep experiment (Figure S1D). The graphical representation of these neurons within the 2D scatter plot enables us to distinguish between MCH putative and OX putative neurons. MCH putative neurons fire maximally during REM sleep and are thought to present a low EMG amplitude and a low average discharge rate of 1.1 ± 0.26 Hz, while OX putative neurons fire maximally during wake and are characterized by a high firing rate, 3.17 ± 0.79 Hz, as both classes of neurons were previously described [18]. Based on these criteria, we assessed MCH putative and OX putative neurons between PWScr^m+/p+^ wild-type and PWScr^m+/p−^ mutant mice. The percentage of recorded MCH putative neurons was unchanged between the two genotypes; although non-statistically significant, a total of 211 OX putative neurons were identified in PWScr^m+/p−^ mutant mice compared with 315 OX putative neurons identified in control mice. This observation suggested that PWS mutants may express a reduced number of OX neurons in the LH (Figure S1D).

### A high proportion of neurons in the LH do not respond to food intake in PWScr^m+/p−^ mutant mice

Mice underwent food deprivation (FD) during the dark period/resting phase (Figure 1B) to increase their drive for food intake. When food was made available again, pellets were automatically provided in the home cages of each mouse (see Methods). The frequency of nose-poke activity to gain food was similar between the two genotypes (PWScr^m+/p−^: 107 trials; PWScr^m+/p+^: 116 trials), indicating no obvious behavioural differences between the two genotypes, as previously described [19, 20].

As for the sleep experiment, our attention fell on the distribution of neuronal modulation between the two genotypes. We found that PWScr^m+/p−^ mutant mice had less than half the number of type II neurons, which have reduced firing during food intake, of control mice (Figure 1G). This reduction may be explained by the fact that mutant mice had twice as many non-responding neurons as control mice. Indeed, type I neurons were almost identical in the two groups (Table S4). These results also suggest for an abnormal distribution of OX neurons in the LH of mutants, as OX neurons drive food intake.

### Sleep homeostasis is disrupted in PWS mutant mice

We explored the 24-h EEG profile of sleep (see Methods) and the responses to sleep loss in a different cohort of animals.

The results confirmed our previous observations that sleep is altered in the PWS mutants [5]. In particular, REM sleep is increased in PWScr^m+/p−^ mutants compared with controls, showing a peak increase at ZT (Zeitgeber Time) 20 and an overall increase in theta power (Figure 2A). These experiments were conducted at standard room temperature of 22°C. Interestingly, the theta profile in mutants did not show circadian rhythm (Figure 2A); PWScr^m+/p−^ mutants had an increase in theta power during the transition between light and dark, when the pressure of sleep is low. Overall, REM sleep abnormalities remained a selective phenotype in these mutants, while the other EEG-determined arousal states were unchanged between the two genotypes (Figure S2A).

**Figure 2.**
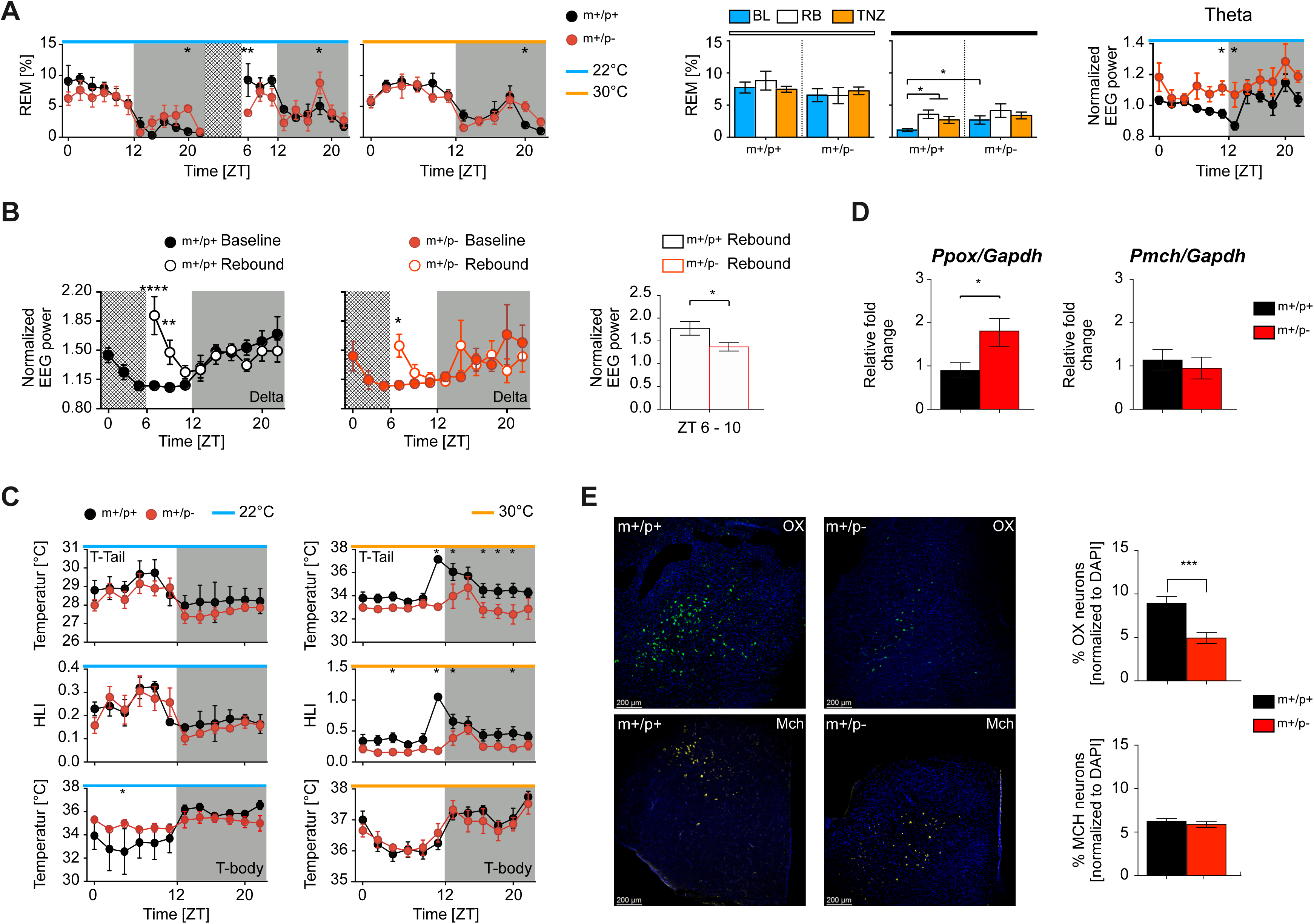
*Snord116* influenced REM sleep and its homeostatic regulation, thermoregulatory response via the orexin system. **A)** Left panel, REM sleep distribution over an uninterrupted 24-h period over a 12-h light/dark cycle as baseline values and the following 18-h after total SD. Recordings were made with animals maintained at 22°C (cyan bar in the graph) or at 30° C (orange bar in the graph). REM sleep was significantly altered in mutant mice relative to control mice at baseline and over the recovery period following 6-h of SD (two-way ANOVA: F (20, 160) = 8.66 p= <0.0001; “time”; F (8, 160) = 3.33; p= <0.001; “genotypes”). REM sleep was also altered in mutant mice relative to control mice when the sleep-wake cycle was recorded at the TNZ (two-way ANOVA: F (11, 88) = 11.61 p= <0.0001; “time”). Data are reported as the percentage in 2 h bins, averaged within genotypes (mean ± SEM; n = 5 PWScr^m+/p+^ and n = 5 PWScr^m+/p−^ mice). PWScr^m+/p−^ mice are represented by red circles, while control mice are represented by black circles. Middle panel, the cumulative amount (mean ± SEM) of REM sleep during the 12-h of the light period (white bar above the bars) and during the 12-h of the dark period (black bar above the bars) for the three groups investigated (recordings at 22°C are shown in blue, after 6-h of SD in white and at 30°C in orange). The within-group statistical analysis was performed by one-way ANOVA, followed by a post hoc analysis with the Bonferroni multiple comparison test. During the dark period, PWScr^m+/p−^ mice showed an increase in REM sleep (F(1.85, 4.74)= 18.06, p = .0082). Student’s unpaired t-test between genotypes indicates an increase in REM sleep in PWScr^m+/p−^ mutant mice relative to PWScr^m+/p+^ mice (t(8) = 2.30, p = .04). The right panel shows the spectral analysis. Theta power at 22°C between genotypes. The theta power of each subject was normalized by the mean of the last 4-h of the light period. Theta power during REM sleep was increased in PWScr^m+/p−^ mice (two-way ANOVA:, F(11,88)= 3.08, p = <.001 “time”; F(1,8)= 8.04, p= .02 “genotypes”). Data are reported in 2-h bins and are shown as the mean ± SEM. **B)** The panel shows the delta power at 22°C between baseline vs rebound for PWScr^m+/p+^ and PWScr^m+/p−^ mice. The delta power of each subject was normalized by the mean of the last 4-h of the light period. Wild-type mice displayed a significant increase in delta power following SD from ZT 6 to ZT 10 (two-way ANOVA: F(11,88)= 28,77 p= <0.0001 “interaction”), while mutant mice showed a mild increase only at ZT 6 (two-way ANOVA: F(11,88)= 12,25 p= <0.0001 “interaction”). Data are reported in 2-h bins and are shown as the mean ± SEM. Right panel, the average of the first 4-h of recovery after SD between the two genotypes for the delta power. PWScr^m+/p−^ mice showed lower delta than PWScr^m+/p+^ mice (unpaired t-test: t(8) = 2.31, p = .04). Values are reported as the mean ± SEM. Asterisks (*) indicate a significant difference between genotypes: * P ≤ .05; **p ≤ .01; *** p ≤ .001; **** p ≤ .0001. Two genotypes were investigated: PWScr^m+/p−^ mice (n= 10, 5 mice at 22°C and 5 mice at 30°C) and PWScr^m+/p+^ mice (n= 10, 5 mice at 22°C and 5 mice at 30°C). **C)** Temperature profile for PWScr^m+/p+^ and PWScr^m+/p−^. The top panel shows the T-tail profile recorded with an infrared thermocamera over 24-h. Recordings were performed under two different environmental conditions: at 22°C (cyan bar over the graph, shown on the left) and at 30°C, corresponding to the thermoneutrality zone (TNZ) (orange bar over the graph, shown on the right). Middle panel, the heat loss index (HLI) was calculated according to the following formula: 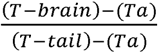 (see Methods) and assessed at 22°C (left) and at the TNZ (right). Bottom panel, the core body temperature profile recorded simultaneously with EEG over 24-h was recorded using a telemetric system in which the probe was subcutaneously implanted. PWScr^m+/p−^ mice are represented by red circles, while PWScr^m+/p+^ mice are represented by black circles. Values are expressed as a 2-h mean ± SEM. At 22°C, no differences were observed between the two genotypes for T-tail and HLI. PWScr^m+/p−^ mice showed an increase in body temperature during the light period at ZT 6 at 22°C (two-way ANOVA: F(11,88)= 3.53, p = .0004 “time”; F(11,88)= 7.86, p = <.0001 “genotypes”). At 30°C, T-tail (two-way ANOVA: F(11,88)= 2.68, p = <.0001; “interaction”) and HLI (two-way ANOVA: main effect of time-of-day, F(11,88)= 4.72, p = <.0001 “interaction”) were increased in PWScr^m+/p+^ mice relative to mutant mice. No differences in body temperature were observed at 30°C. Asterisks (*) indicate a significant difference between genotypes (* P ≤ .05). Two genotypes were investigated: PWScr^m+/p−^ mice (n= 10, 5 mice at 22°C and 5 mice at 30°C) and PWScr^m+/p+^ mice (n= 10, 5 mice at 22°C and 5 mice at 30°C). **D)** *Ppox* and *Pmch* gene expression analysis in PWScr^m+/p−^ mice versus controls. PWScr^m+/p−^ mice (red bar) showed an increase in *Ppox* (unpaired t-test: t(8) = 2.49, p = .03) compared with PWScr^m+/p+^ mice (black bar). No differences in the mRNA *Pmch* level was observed between the two genotypes. Values are reported as the mean ± SEM. Asterisks (*) indicate a significant difference between genotypes: * P ≤ .05. **E)** Cell count distribution of orexin (OX) immunoreactive neurons (upper) and melanin concentrate hormone (MCH) immunoreactive neurons (below) in the lateral hypothalamus of PWScr^m+/p−^ mice (right) versus controls (left). Coronal sections were stained with OX- and MCH-specific antibodies, counterstained with DAPI and scored. Values are expressed as the percentage of positive neurons relative to all stained nuclei (mean ± SEM). The number of OX^+^ neurons was reduced in the PWScr^m+/p−^ mouse group (unpaired t-test: t(33) = 3.85, p = .0005). No differences were found in the number of MCH^+^ neurons. Asterisks (*) indicate a significant difference between genotypes: *** p ≤ .001. Two genotypes were investigated: PWScr^m+/p−^ mice (n=4) and PWScr^m+/p+^ mice (n= 4).

Moreover, we tested for the first time the homeostatic EEG response following SD in PWScr^m+/p−^ mutant mice compared with littermate controls. Compared with control mice, PWScr^m+/p−^ mice showed reduced delta activity during non-REM (NREM) sleep rebound following SD (Figure 2B), although the total amount of NREM sleep was unaltered over the long-term recovery process (Figure S2A). Notably, delta power is the best electrophysiological marker of sleep propensity, although the temporal distribution of its rebound following SD is less characterised. The behaviour of delta sleep in mutant mice suggests a dysregulation in the daily distribution of sleep pressure in these mice.

Our results also showed a significant decrease in REM sleep during the first 2-h of rebound in mutants compared with controls (Figure 2A), confirming both the alteration of homeostatic control in mutants and the alteration of REM sleep.

### Thermoneutrality does not increase REM sleep in mutants

REM sleep is an evolutionarily recent physiological state of sleep that, in mammals, is largely dependent on the environmental temperature [21, 22]. The maximum REM sleep expression in mice is attained when the environmental temperature is near 29°C. The latter represents a thermoneutral zone (TNZ) for this species [23].

In a new set of experiments, we modified the temperature environment of the animals to allow the duration of REM sleep to reach a maximum [23]. Mice were housed for 5 weeks at 30°C, and then, EEG/EMG were recorded for 24-h at the TNZ. Mutant mice showed a significant increase in all sleep stages (Figure S2) compared with wild-type mice, but REM sleep did not significantly increase in the mutant mice, perhaps due to the high REM sleep percentage at baseline (Figures 2A, S2C and S2D). However, PWScr^m+/p−^ mutant mice maintained a significant increase in REM sleep compared with controls at ZT 20 (Figures 2A and S2A).

### Peripheral thermoregulatory responses are absent in PWS mutants

We tested whether loss of paternally expressed *Snord116* affects peripheral responses and whether these changes impact the overall body temperature and body weight of the animals. We monitored the peripheral and cutaneous body temperature at the head (T-head) and tail (T-tail), and consequently, we derived the heat loss index (HLI) (see Methods). Recordings were made at room temperatures of both 22°C and 29-30°C (TNZ). The TNZ imposes body temperature adjustments by changing the vasomotor tone in specialized heat exchange organs, such as the tail, in mice [24]. We observed an increase in the vasculature tail skin tone at the TNZ in both genotypes compared with 22°C (Figure 2C, upper panel). However, greater tail vasodilatation was observed in control mice, particularly between the switch from light to dark periods. These results indicate that while at TNZ, wild-type control mice present a proper peripheral thermoregulatory response when the pressure of sleep is low, which is instrumental in maintaining the core body temperature of the animal at this time of the day, mutant mice lack such an important physiological response. A significant increase in the HLI was recorded in both genotypes, but a greater increase was found in PWScr^m+/p+^ wild-type mice than in mutant mice (Figure 2C, middle panel). We also observed that the peripheral body temperature recorded at 22°C increased in PWScr^m+/p−^ mutant mice compared with control mice during the light period (Figure 2C, bottom panel), suggesting a lack of homeostatic control when sleep is physiologically facilitated in mice. Moreover, mutant mice failed to show the circadian oscillatory profile of temperature at RT that is present in PWScr^m+/p+^ mice. However, at TNZ, the difference between the two genotypes during the light period was no longer observed. Furthermore, in agreement with previous reports [20], we observed that PWScr^m+/p−^ mutant mice had a smaller body size when kept at 22°C (Figure S3). Interestingly, the smaller body size was observed both when the mutant mice were exposed to 30°C later in their development (e.g., for only 5 weeks) and when they were exposed to the TNZ from birth and kept at the TNZ for up to 20 weeks of age (Figure S3). However, while the former 5-week experimental strategy resulted in an increase in body weight in both cohorts of animals, which was more pronounced in wild-type animals than in mutants, the latter (birth) strategy resulted in no changes over time according to temperature (Figure S3). Notably, maintaining at the TNZ from birth resulted in an overall body weight in mutants that was comparable to the body weight at RT. The latter observation suggests that the TNZ is a condition that permits an increase in body weight in PWS animals, although this cannot neutralise the differences with wild-type animals under the same conditions.

### Lack of *Snord116* impairs the OX system in the LH

To assess the regulation of the main neuropeptides in the LH, we examined the expression of OX and MCH. Thus, we assessed the precursor of MCH (*Pmch*) and the prepro-OX (*Ppox*). We examined three conditions (Figure S4A): at ZT 6, we tested both the baseline condition (T0) and the effects of SD (T1); and at ZT 7, 1-h after SD (T2), we tested the effects during the recovery from the loss of sleep.

We observed that PWScr^m+/p−^ mutant mice showed a significant increase in *Ppox* at T0 baseline compared with control mice, while *Pmch* was unchanged in both genotypes at T0 (Figure 2D). At T1, both *Ppox* and *Pmch* were significantly reduced in PWScr^m+/p−^ mice compared with control mice (Figure S4B), confirming that the homeostatic response in mutants is reduced and that these two systems are dysregulated in mutant mice. At T2, the difference between the two genotypes for *Ppox* and *Pmch* was repristinated (Figure S4B), confirming that the effects of SD in mutants are only acute and follow the immediate response after sleep deprivation.

Then, we evaluated the receptors. We assessed mRNA levels of OX receptor-1 (*Ox1R*) OX receptor-2 (*Ox2R*) and MCH receptor 1 (*Mch1R*) in the hypothalamus and in other brain areas as control measures. The expression levels were overall unchanged between the two genotypes, except for *Ox2R* in the parietal cortex (Figure S4C).

The overall contrast between neuropeptides and the mRNA levels of their receptors prompted us to quantify whether these chemical changes within the OX and MCH systems had consequences for the organization of the neuronal populations of the LH. We observed a significant reduction in OX neurons in the LH of PWScr^m+/p−^ mutant mice (PWScr^m+/p+^ mice 95 □±□9; PWScr^m+/p−^ mice 53 □±□10; p= .0005; Figure 2E), while MCH neurons, which are located near the OX neurons, were unaffected (PWScr^m+/p+^ mice 66 □±□5; PWScr^m+/p−^ mice 63 □±□5; Figure 2E). The discrepancy between the increased *Ppox* and the reduced neuronal population may be due to a mechanism to compensate for the low number of neurons with an overproduction of the peptide.

### Loss of *Snord116* leads to transcriptional reprogramming in the hypothalamus

Next, we investigated whether the loss of *Snord116* affects overall gene expression in the hypothalamus of PWScr^m+/p−^ mutant mice compared with PWScr^m+/p+^ control mice by performing RNA sequencing (RNA-seq). The mice were investigated at the beginning of the light period ZT 0 (group 1, G1), 6-h later at ZT 6 (group 2, G2) and at ZT 6 but following 6-h of SD (group 3, G3). First, we sought to identify the differentially expressed genes (DEGs) between the two genotypes in the G1 group. We identified 4777 downregulated genes and 4271 upregulated genes between genotypes (adjusted p < 0.05; see Methods) and a minimum fold change of 2 (Table S8). Next, we compared the down- and upregulated DEGs (Figure 3 A-B) in G1 PWScr^m+/p−^ mutant mice with DEGs derived from post-mortem hypothalamic data from PWS patients vs normal patients [25].

**Figure 3.**
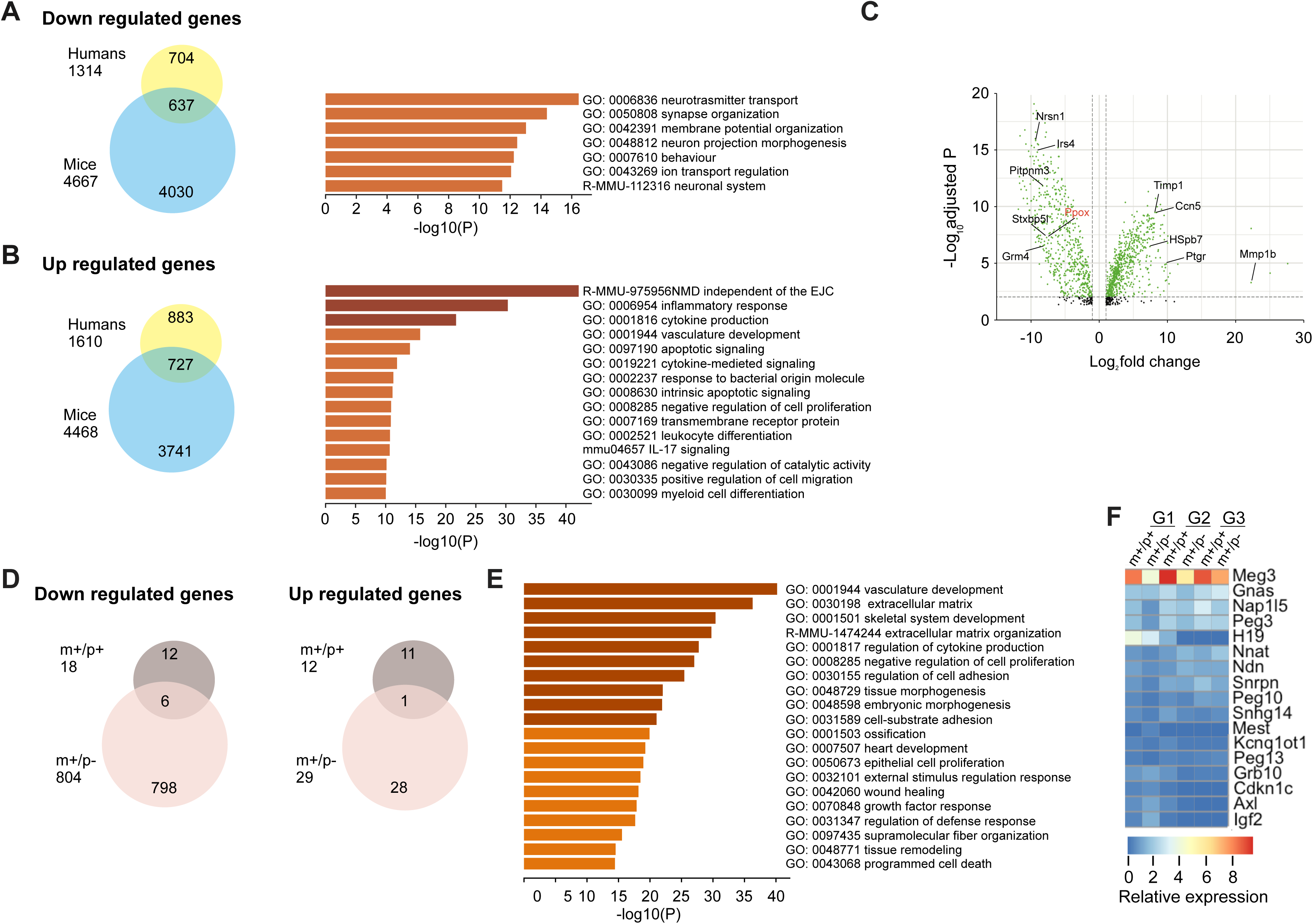
*Snord116* loss significantly impacts molecular machinery in the hypothalamus. **A-B)** Venn diagrams illustrating the number of differentially expressed genes (DEGs) that are down- (A) and upregulated (B) in the hypothalamus of Prader–Willi syndrome (PWS) PWScr^m+/p−^ mice relative to control mice and that overlap in human patients (according to [25]. The results of gene ontology (GO) enrichment analysis of biological processes for the overlapping DEGs are also shown in both A and B. **C)** Volcano plots of 637 and 727 DEGs in PWScr^m+/p−^ mice in group 1 (G1; non-sleep deprived). **D)** Significantly down- and upregulated genes in the hypothalamus of PWScr^m+/p−^ mutant mice compared with PWScr^m+/p+^ control mice affected by sleep deprivation (G2 vs G3). **E)** GO enrichment analysis of biological processes for 833 (804 down- and 29 upregulated genes in panel D) DEGs in PWScr^m+/p−^mutant mice that are significantly affected by sleep deprivation. **F)** Heatmap of the relative expression of imprinted genes common in humans and mice assessed in PWScr^m+/p−^ mutant mice compared with the PWScr^m+/p+^ mice in G1, G2 and G3. See also Tables S8 and S9.

More than 40% of human DEGs overlapped with mouse DEGs, suggesting that loss of *Snord116* recapitulates approximately 40% of the transcriptional changes found in the human hypothalamus. To more closely evaluate the functional importance of these overlapping DEGs, we performed gene ontology (GO) enrichment analysis using Metascape [26]. We found enrichment of several biological processes important for normal neural functions among the downregulated DEG genes, while inflammatory systems were the processes significantly enriched among the upregulated DEGs (Figure 3A-B).

Interestingly, we found that *Ppox* was significantly downregulated in PWScr^m+/p−^ mutant mice and in human PWS patients (Figure 3C), confirming that OX is involved in the abnormal regulation of REM sleep in PWS.

Next, we focused on how SD affects gene expression in the hypothalamus in both normal and PWScr^m+/p−^ mutant mice. We compared DEGs in G2 versus G3 of each genotype (Figure 3D). We found that 6 h of SD largely influenced the transcriptome of mutant mice (i.e., 833 DEGs) compared with control mice (i.e., 30 DEGs). Surprisingly, in the mutant dataset, most of the DEGs were downregulated (804 genes downregulated vs 29 upregulated). GO analysis of all 833 DEGs revealed enrichment of cell organization, development and growth-related processes (Figure 3E), indicating that poor sleep negatively impacts the hypothalamus in PWS subjects. Finally, we investigated the RNA-seq dataset to determine whether *Snord116* loss significantly affects the expression of imprinted genes in humans and mice (refer to Table S9). Maternally expressed imprinted genes, such as *Meg3*, *Gnas*, and *H19*, as well as paternally expressed imprinted genes, such as *Nap1l5* and *Peg3*, were differentially expressed in mutant mice before and after SD (Figure 3F).

### OX regulation in the hypothalamus depends on two paternally expressed genes, *Snord116* and *Peg3*

We next sought to confirm selected physiologically relevant DEGs identified within the RNA sequencing dataset by qRT-PCR. Among maternally and paternally imprinted genes that were altered in the hypothalamus of PWScr^m+/p−^ mutant mice, *Peg3* was the only confirmed gene, showing a remarkable increase in expression in PWScr^m+/p−^ mice (Figure 4A) compared with the control mice (Figure S5A). Peg3 binds to DNA based on its multiple zinc finger motifs and nuclear localization [27, 28]. Interestingly, a recent study found an association between *Peg3* and OX expression [29]. However, whether Peg3 is able to bind and regulate the expression of OX in living mice remains unclear. We noticed that two conserved promoter regions within a 3.2-kb fragment located upstream of the *Ppox* gene (Figure S5F) target specific expression within the LH [30]. We showed that PEG3 binds one of these regions. Specifically, chromatin immunoprecipitation analysis (ChIP) [31] followed by quantitative real-time PCR (ChIP qRT-PCR, and Figure 4A) revealed that PEG3 binds the 2.5 kb OX regulatory element region in the hypothalamic mouse brain. Interestingly, a significant reduction in PEG3 binding was observed in PWScr^m+/p−^ mice (Figure 4A), which may be associated with a significant reduction in OX-expressing neurons in mutant mice. Our results show that PEG3 positively regulates the expression of *Ppox* by enhancing its expression, as indicated by the presence of a strong enrichment of the H3K4me2/3 in the same region (Figure 4A). Conversely, H3K27me3, which is a marker of condensed chromatin[32], showed no enrichment in either genotype. The GAPDH promoter was used as a negative control for the ChIP experiment (Figure S5E). These data reinforce the evidence that PEG3 contributes to the regulation of *Ppox* by enhancing its expression (Figure 4A).

**Figure 4.**
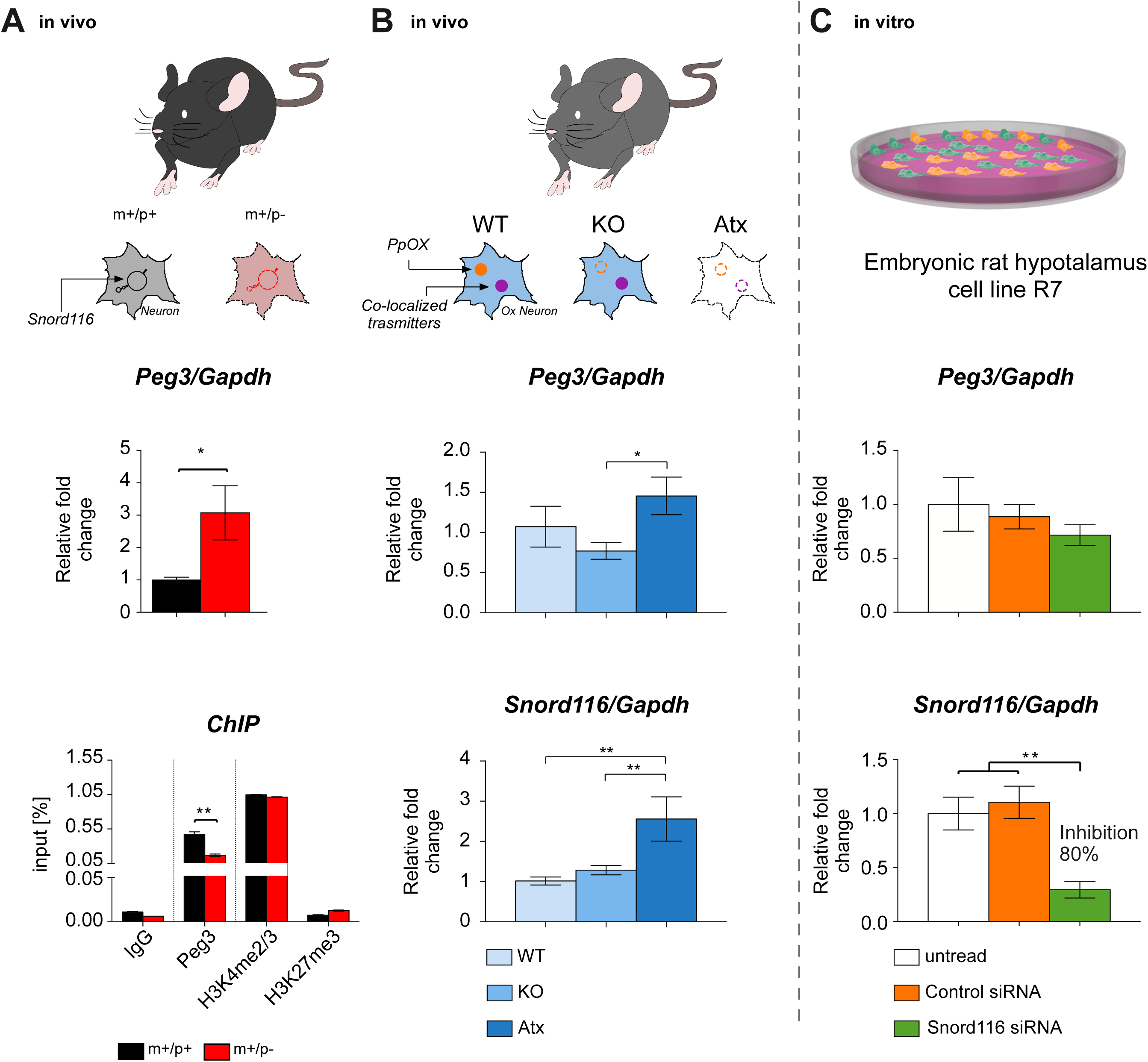
*Snord116* and *Peg3* play roles in the formation and maintenance of OX neurons. *Peg3* regulates orexin expression in an independent manner from paternal *Snord116*. **A)** Upper panel, the gene expression analysis of *Peg3* in PWScr^m+/p−^ mice (red) versus controls (black). *Peg3* mRNA assessed by qRT-PCR was significantly increased in PWScr^m+/p−^ mice compared with PWScr^m+/p+^ mice (unpaired t-test: t(6) = 2.45, p = .04). Values expressed are relative to the wild-type control mean ± SEM. *Gapdh* was used as a housekeeping gene; see Methods. Asterisks (*) indicate a significant difference between genotypes. Bottom panel, ChIP analysis of PEG3 binding to the *Ppox* promoter region in PWScr^m+/p−^ mice (red) versus controls (black). PEG3 binding was lower in PWScr^m+/p−^ mice than in PWScr^m+/p+^ mice (unpaired t-test: t(2) = 7.11, p = .01). Values are expressed as the mean of the input ± standard deviation. Asterisks (*) indicate a significant difference between genotypes. **B)** *Peg3* gene expression (upper panel) and *Snord116* gene expression (bottom panel) in *Ppox* knockout (KO) and orexin neuron-ablated (ataxin-3 [Atx] mice). One-way ANOVA indicated that *Peg3* was significantly increased in Atx mice relative to KO mice, (F(2,16)= 0.02, Bonferroni post hoc test p = .03). Snord116 was increased in Atx mice relative to KO and control mice (WT) (one-way ANOVA F(2,16)= 3.50, Bonferroni post hoc test p = .002). Values are reported as the mean ± SEM. Asterisks (*) indicate a significant difference between genotypes. The following genotypes of narcoleptic mice were investigated: WT (n=4), KO (n= 12) and Atx (n=4). **C)** *Snord116* and *Peg3* gene expression analysis in the *Snord116*-siRNA-treated immortalised hypothalamic rat cell line. *Snord116*-siRNA (green bars) reduced the expression of the *Snord116* gene compared with untreated cells or scrambled siRNA-treated cells (white and orange bars) (one-way ANOVA: F(2, 6)= 11.36, Bonferroni post hoc test p = .009). *Peg3* mRNA levels were unchanged, similar to *Snord116*-SiRNA, in untreated cells and scrambled siRNA-treated cells. The experiment was conducted in triplicate. Data are presented as the mean ± SEM. Asterisks (*) indicate a significant difference between genotypes: * P ≤ .05; **p ≤ .01.

Then, we sought to investigate whether *Peg3* or *Snord116* plays any causal role in the regulation and formation of OX neurons, and we assessed the expression of these two genes in OX-deficient mice. Specifically, we used two genetic mouse models of OX deficiency, *Ppox* knockout (KO) mice [9] and OX neuron-ablated (ataxin-3 [Atx]) mice [33] (Figure 4B).

Atx mice selectively degenerate postnatally, and a loss of 99% of neurons or more occurs by the age of 4 months [33]. Therefore, in Atx mice, not only are OX neurons ablated, but co-transmitters are also eliminated [34]. Conversely, KO mice are only deficient in *Ppox* (Figure 4B).

Interestingly, we observed that *Peg3* was significantly increased in Atx mice compared with KO mice (p= .03). Moreover, *Snord116* was also increased in Atx mice compared with KO mice and control mice (p= .03; Figure 4B). Other imprinted genes tested were unchanged between genotypes (Figure S5B). These data suggest that dysregulation of both *Snord116* and *Peg3* are important and associated with the physiological maintenance of OX neurons. Indeed, we observed that both genes were significantly altered only in narcoleptic mice, in which OX neurons and their co-transmitters were ablated, but not in narcoleptic mice lacking only Ppox.

Finally, to test whether the alteration of *Peg3* observed in the PWScr^m+/p−^ mice and in the Atx mice is linked to the absence of *Snord116* and whether *Snord116* modulates the expression of *Peg3*, we used an RNA-interfering approach aimed at silencing *Snord116* in immortalised embryonic rat hypothalamus cells (embryonic rat hypothalamus cell line R7). Specifically, cells transfected with a *Snord116*-specific siRNA showed significant *Snord116* inhibition of approximately 80% at 48 h after transfection (Figure 4C). However, the levels of *Peg3* were unchanged (Figure 4C), indicating that *Snord116* does not regulate the expression of the *Peg3* gene. *Pmch* and *Snurp* were also unchanged (Figure S5).

## Discussion

Overall, these new results reinforce the evidence that loss of *Snord116* plays a crucial role in the regulation of REM sleep and thermoregulation, both phenotypes being dysregulated in PWScr^m+/p−^ mutant mice and patients [5, 35]. In particular, we repeatedly observed REM sleep alterations at a particular time of the day when the pressure of sleep was increased [36, 37]; in mice, this occurs at a time in the sleep-wake cycle that corresponds to the “*siesta time*” [36, 37]. This new evidence could be instrumental for future experimental designs aimed at addressing temporally precise pharmacological treatments.

*Snord116* is one of the most important candidate genes involved in the aetiology of the major endophenotypes of PWS. Individuals lacking the *SNORD116* snoRNA cluster and the *IPW* (Imprinted In Prader–Willi Syndrome) gene suffer the same characteristics of failure to thrive, hypotonia, and hyperphagia that are observed in subjects with larger deletions and maternal uniparental disomy [38, 39].

Our *Snord116* mouse model remarkably recapitulates all the sleep deficits of human PWS, but only incompletely mimics the metabolic alterations. Indeed, both PWS mice and human patients show an alteration of body weight, but the two species show opposite phenotypes; indeed, PWS patients exhibit increased body weight, while mouse models across different studies show reduced weight [20, 40]. This overall result suggests that evolutionary divergency between the two species may have played an important role in the metabolism of PWS, but to date, there is no evidence for such a change.

We report for the first time a lack of neuromodulation of the LH in *Snord116* mutant mice, which is accompanied by an imbalance between OX and MCH neurons, causing a 60% reduction in OX neurons in the LH. Within the same 15q11-q13 region, it was previously reported that *Magel2* KO mice lose 40% of OX neurons [41]. These results suggest that multiple paternally expressed genes within the PWS region regulate the OX system in the LH, most likely in a dose-dependent manner. At baseline, we observed that fire discharge associated with sleep (*i.e.*, S-max neurons) is higher in mutant mice than in controls, suggesting that the role of *Snord116* that originates from the LH is important in the determination of abnormal sleep in PWS. Indeed, several lines of evidence indicate that LH exerts pivotal control of cortical sleep, including REM sleep [42]. Furthermore, while we observed that the proportion of S-max and W-max neurons significantly changed after 6 h of SD with an increase in sleep neurons of approximately 13% in wild-type control mice, PWS mutants lack this neuronal response by the LH to sleep loss. In the LH, OX and MCH systems exert opposing effects on REM sleep [43]: OX suppresses while MCH promotes REM sleep. Therefore, the imbalance between OX and MCH systems in PWS mutants is in agreement with an increase in REM sleep.

Additionally, OX and MCH neurons in the LH regulate food intake and metabolism [44, 45]. We found that mutant mice displayed a higher proportion of neurons, which were classified as non-responding neurons (type III), relative to food intake than controls. In contrast, type II neurons, which are downregulated in the LH during eating, were significantly reduced in mutants. It has been demonstrated that OX neurons decrease their firing during eating and remain in a depressed state throughout the entire eating phase [46]. These findings suggest a lack of regulatory feedback mechanisms mediated by the OX system in relation to food intake in the LH of PWS model mice, a phenomenon that can relate to the hyperphagia and obesity phenotype observed in PWS patients. This evidence of neuromodulatory dysregulation of the LH is in agreement with previous results [19], which describe that a late onset of mild hyperphagia and obesity in mice can be induced only when *Snord116* is selectively disrupted in the hypothalamus in adult mice.

Dysregulation of the OX system has been reported in a few clinical studies, although to date, the results remain contradictory. One study described an increase in the OX-A level in PWS subjects [47], while another showed a decrease in the peptide in the cerebrospinal fluid [11]. Taken together, these findings call for an alteration of the OX system in PWS, although the contrasting results may also suggest that the loss of function of paternal alleles within the PWS region results in an increased phenotypic variance.

Our data suggest that *Snord116* is essential for the regulation, formation and maintenance of the OX system in the LH. We assessed *Snord116* expression in two different strains with OX deficiency, OX/ataxin-3 (Atx) mice and OX peptide knockout (KO) mice [9, 33], and we found that only Atx mice showed a significant increase in *Snord116*. This line selectively loses OX neurons and their co-transmitters. In contrast, in KO mice that do not display a loss of OX neurons but are lacking OX peptides, the level of *Snord116* was unaffected. The increase in *Snord116* in mice with depletion of OX neurons is probably a compensatory mechanism. Our conclusions are supported by a recent study [48] that described the role of *Peg3* in the development and expression of OX and MCH neurons. Our study demonstrates for the first time that Peg3 is able to bind OX promotors by increasing the expression of OX. However, a reduced binding of Peg 3 was observed in mutant mice relative to control mice, which may be explained by a reduction in OX neurons. Thus, *Peg3* was selectively altered in our PWScr^m+/p−^ mutant mice as well as in narcoleptic mice. In particular, *Peg3* was found to be altered only in Atx narcoleptic mice, while it was unchanged in Ppox KO mice. This may be explained by the fact that Peg3 binds the OX promoter, which is not altered in KO mice because they have a null mutation. Our study reveals that these two paternally imprinted genes, *Snord116* and *Peg3*, are unlikely to interact with each other, but both contribute to the development and functions of OX neurons. Indeed, in our *in vitro* experiment with immortalised hypothalamic cells, knockdown of *Snord116* did not change the levels of *Peg3*.

All these evidence are in line with the pioneering evidence in androgenetic mice [1] that paternally imprinted genes are important for the formation of the hypothalamus. In particular, our study implies that *Snord116* and *Peg3* plays a crucial role in the formation of OX neurons.

We observed that PWScr^m+/p−^ mutant mice present a high body temperature coupled with an increase and lack of appropriate thermoregulatory responses. Thermoregulation is tightly integrated with the regulation of sleep and is also controlled by OX neuromodulation [34, 49]. For example, narcoleptic subjects exhibit a paradoxical lower core body temperature while awake [50] and a higher body temperature during sleep [51, 52]. In mammals, in physiological conditions, peripheral vasodilatation helps decrease the core body temperature during sleep initiation. Our mutant mice displayed a high body temperature during the light period, which corresponded to their resting phase/subjective sleep. Interestingly, narcoleptic mice show body similar temperature abnormalities [53]. Moreover, when PWS mutant mice were kept at the TNZ, where resting metabolic rate remains stable and where REM sleep is preferentially increased [54, 55], we observed a surprising thermoregulatory response coupled with an altered homeostatic REM sleep response. REM sleep did not increase, and the peripheral thermoregulatory response of the mutant mice resembled what would be expected in a sub-neutrality (e.g., 22°C) environment. REM sleep is a stage of sleep in which thermoregulation is suspended; for this reason, REM sleep expression is more sensitive to ambient temperature manipulation than NREM sleep [54, 55]. A recent study [56] described that endotherms have evolved neural circuits to opportunistically promote REM sleep when the need for thermoregulatory defence is minimized, such as in TNZ conditions, suggesting a tight link between thermoregulation and REM sleep. In our study, the increase in body weight observed at the TNZ suggests that thermoneutrality is a permissive condition that induces body weight gain but does not compensate for the metabolic abnormality in these mice. Indeed, the differences with wild-type mice remained unchanged. Mutants showed growth retardation at both environmental conditions investigated, namely, at both 22°C and 30°C.

The transcriptomic analysis in the hypothalamus suggests that loss of *Snord116* might negatively affect synaptic organization while promoting inflammatory responses in the hypothalamus, as previously observed in post-mortem hypothalamic brain tissue from PWS patients [25]. GO analysis of all 833 DEGs in the mutant hypothalamus following SD suggested, instead, that loss of *Snord116* leads to a homeostatic response that relies on several cellular growth processes of the hypothalamus (Figure 3E). This result indicates that defects in sleep homeostasis in PWS can be derived from development processes of the hypothalamus, as has been described in other neurodevelopmental disorders [57–59].

In conclusion, our study demonstrates for the first time that paternally expressed genetic elements in the LH affect the dynamics between neuronal activity in the LH and cortical EEG sleep states. This new evidence reinforces the recent hypothesis that genomic imprinting plays a crucial role in mammalian sleep [3] and confirms original studies in androgenetic chimeric mice [1], suggesting that the paternal genome may account for regulatory mechanisms in the hypothalamus.

## Methods

### Animal husbandry, breeding, and genotyping

All animals were housed in controlled temperature conditions (22 ± 1 °C) under a 12 h light/dark cycle (light on 08:00–20:00), with libitum access to food (standard chow diet) and water unless otherwise required by the experimental procedure.

Experiments were performed using adult mice at 15-17 weeks with paternal deletion of the *Snord116* gene and *IPW* exons A1/A2 B [16] (PWScr^m+/p−^) and their wild-type littermates as control mice (PWScr^m+/p+^). To maintain the colony, mice were bred and kept through paternal inheritance on a C57BL/6J background. Genotyping was performed as previously described [5]. Briefly, PCR analysis of genomic DNA from ear punches was performed using the primer pair PWScrF1/PWScrR2 (5’-AGAATCGCTTGAACCCAGGA and 5’-GAGAAGCCCTGTAACATGTCA, respectively). The deletion of PWScr resulted in a PCR product of approximately 300 bp, which was absent in the wild-type genotype.

For experiments in narcoleptic mice, 3 age-matched groups of male and female congenic mice from two different strains (≥ 9 generations of backcrossing) on the C57Bl/6J genetic background were used, as follows: (i) mice with congenital deficiency of OX gene (*Ppox*) knockout (KO) [9] (n=11); (ii) mice hemizygous for a transgene (ataxin-3 [Atx] mice), the targeted expression of which causes selective ablation of OX neurons [33] (n=4); and (iii) wild-type controls (n= 4). Mouse colonies were maintained in the facilities of the Department of Biomedical and NeuroMotor Sciences at the University of Bologna, Italy. Mice were housed under a 12-h light/dark cycle at an ambient temperature of 25°C with free access to water and food (4RF21 diet, Mucedola, Settimo Milanese, Italy).

All animal procedures were approved by the Animal Research Committee and the Veterinary Office of Italy for Istituto Italiano di Tecnologia (IIT) Genova. All efforts were made to minimize the number of animals used and any pain and discomfort according to the principles of the 3Rs [60].

### Experimental approach

#### Neuronal activity in the mouse LH is time-locked with different behavioural and physiological states of the brain

To identify neuronal populations in the LH that control different physiological behaviours, we adopted a combined experimental approach (Figure 1A) in which we monitored single unit activity (SUA) at the LH in parallel with the cortical state of each animal by means of electroencephalogram and electromyogram (EEG/EMG) recordings. The EEG/EMG information allows the identification of wakefulness, REM sleep and NREM sleep states in the animal, which represent the three main behavioural states of the mammalian brain. Mice were kept throughout a regular sleep-wake cycle, and EEG/EMG/SUA were recorded during the light-to-dark transition (baseline 1, B1) and during the dark-to-light transition (baseline 2, B2) to test the neuronal responses during the minimum and maximum levels of sleep pressure, respectively (Figure 1B upper panel). Then, each mouse underwent sleep deprivation (SD) to test the homeostatic response following sleep loss (Figure 1B middle panel). A total of 569 well-sorted units were successfully recorded from PWScr^m+/p+^ wild-type mice, and 548 units were recorded from PWScr^m+/p−^ mutant mice across all time points.

Based on the distribution of the firing activity of each unit, we were able to distinguish populations of putative neurons in the LH that were time-locked with the occurrence of specific sleep-wake states (Figures 1C and 1E). In particular, we identified neurons (Figure 1E) associated with sleep (S-max) or wakefulness (W-max). However, we distinguished between NREM (NR-max), REM (R-max) neurons and neurons firing in both stages (NRR-max) (Table S1). We also identified a group of neurons that did not show significant changes in their firing rate across the sleep-wake states, and we called this latter class of neurons “ws” (wake and sleep).

Moreover, to test the other fundamental function of LH, feeding, we food deprived each animal during the 12 h of darkness of the light/dark cycle, and then, when food was made available, we evaluated the neuronal responses to food intake (Figure 1B lower panel). In this experiment, a total of 147 units were identified from PWScr^m+/p+^ mice and 74 units from the PWScr^m+/p−^ mice in response to food deprivation.

Following the food deprivation protocol, we also distinguished three different classes of neurons in the LH that respond to food intake. In particular, we defined type I and II neurons as those that fire maximally after or before nose-poking/feeding activity, respectively (Figures 1D and 1F). In this experiment, a third class of neurons, type III neurons, was defined as non-responding neurons.

### Experimental protocols

#### LH neuronal activity recordings

To investigate the neuronal dynamics of the lateral hypothalamus (LH), EEG/EMG and SUA were simultaneously recorded in eight male adult PWScr^m+/p−^ and PWScr^m+/p+^ mice. Animals were individually housed after surgery in their home cages with a food hopper (a hole with an infrared beam) connected to a food dispenser that automatically delivers food pellets after a nose poke (powered by AM-Microsystems [61]. The food used was 20 mg dustless precision pellets (Bioserv). After a 7-day recovery period after surgery, each mouse was connected to a flexible cable and swivel that allowed free movement within the cages; the mice were habituated for 2 days to the cable before SUA and EEG/EMG recordings (Figure 1A). Recordings of EEG/EMG with SUA were acquired at two different baseline (BL) time points (B1 and B2; Figure 1A) for 2 h according to the C Process and S Process of sleep. Next, to investigate sleep homeostasis, SUA and EEG/EMG were recorded during the first hour of the rebound period (RB, Figure 1A) following 6 h of total sleep deprivation (SD) and at other two time points over the 18-h recovery period (SD1 and SD2, Figure 1A).

After a restoration period of 5 days from the previous SD, animals were fasted for 12 h during the dark period, which corresponds to the active phase of mice. At the beginning of the light period, mice were fed. A few food pellets were initially provided to encourage the mice to eat, and thereafter, food pellets were provided only after nose-poke activity. SUA was continuously recorded for 2 h after being fed (Figure 1A). EEG/EMG was not recorded during this time.

#### Homeostatic investigation of sleep

To investigate the role of the *Snord116* gene in the regulation of the homeostatic component of sleep, we recorded EEG/EMG in ten male adult PWScr^m+/p−^ and PWScr^m+/p+^ mice over 24 h of baseline (BL) at 22 ± 1°C. The BL recordings began at ZT 0 (the time of light onset in the 12 h light/dark cycle), and then, mice were sleep deprived during the first 6 h of the light phase (ZT 0–6) by gentle handling (introduction of novel objects into the cage, tapping on the cage, and, when necessary, delicately touching) and then allowed 18 h of recovery (ZT 6–24, RB).

#### Homeostatic investigation of REM sleep

To investigate the REM sleep propensity, we housed male adult PWScr^m+/p−^ and PWScr^m+/p+^ mice for five weeks at 30 ± 1°C, which is close to or within the thermoneutral zone (TNZ) of mice [62, 63]. Then, these mice underwent continuous recordings for 24 h at 30°C ambient temperature while undisturbed and freely moving in their cages.

#### Surgery

Mice were anaesthetised using 1.5%–2.5% isoflurane in oxygen and placed in a stereotaxic frame (David Kopf Instruments, Tujunga, CA). To assess the sleep-wake cycle, mice were surgically implanted with a telemetric transmitter (volume, 1.9 cm 3; total weight, 3.9 g; TL11M2-F20-EET; DSI, St. Paul, MN, USA) connected to electrodes for continuous EEG/EMG recordings. A wireless EEG transmitter/receiver, which also contained a sensor to detect body temperature, was subcutaneously implanted. Specifically, EEG wire electrodes were implanted into the frontal cortex (coordinates: 2 mm posterior to the bregma and 2 mm lateral to the midline in the right parietal skull) and the parietal cortex (coordinates: 3 mm anterior to the lambda and 2 mm lateral to the midline in the right frontal skull). EMG was recorded by two stainless steel wires inserted bilaterally into the neck muscles. Subsequently, to record SUA, a tetrode-based 16-channel micro-wire array (4 × 4, 5.3 mm, 100-200-1250) of silicon probes (NeuroNexus Technologies) was implanted into the LH (coordinates relative to the bregma: –1.45 mm anteroposterior, –1.0 mm mediolateral, and –4.8 mm dorsoventral) of the contralateral hemisphere with respect to the EEG electrodes (Figure 1A). Mice were operated on by performing a small craniotomy, and the dura mater was removed for placement of the tetrodes in the LH. Two screws (1 mm diameter) were used to anchor the implant. One screw placed in the cerebellum was wrapped by the electrode ground. Next, the tetrode and the ground wire were covered with dental acrylic. Following surgery, all animals were administered paracetamol (200 mg/kg; once a day; PO; Tempra) and enrofloxacine (10 mg/kg; once a day; SC; Baytril) for two days.

#### Histology

At the end of the last recording session, animals were sacrificed, and the locations of the recording electrodes were verified histologically. Mice were anaesthetised using pentobarbital anaesthesia (100 mgr/kg, i.p.), and microlesions were made at the tip of one or two microwires by passing a small current (5 m A, 10 s). The animals were transcardially perfused with 10 ml of phosphate-buffered saline (PBS) before infusion of 4% paraformaldehyde (PFA) in PBS. The brains were removed and equilibrated in 30% sucrose, sectioned at 40 µm on a freezing microtome, mounted onto gelatin-coated slides, air dried, dehydrated in ethanol, stained with Nissl substance, cleared with xylene, and cover slipped with DPX (Figure 1A). Three mice were excluded from the study because the electrode was not correctly placed in the LH (Figure 1A).

#### Thermal imaging

Surface body temperature was continuously recorded in PWScr^m+/p−^ and PWScr^m+/p+^ mice for 24 h at 22°C and 30°C using an infrared thermocamera (FLIR A20). The thermocamera was positioned above the cage where the mice were individually housed. Head (T-head) and tail temperatures (T-tail) were obtained by manually analysing the recorded video (1 frame/s) using a dedicated software program (Thermocam Research, FLIR). T-head was used to compute the heat loss index (HLI). HLI was calculated by the equation (T-head - Ta)/(T-tail - Ta) [71]. T-head, T-tail and HLI values were collapsed into 2-h bins and compared between conditions, before and after drug treatment, in the two experimental groups.

Additionally, body weight was measured weekly in small groups of both PWScr^m+/p−^ and PWScr^m+/p+^ mice for 5 weeks at 22°C and at the TNZ (n= 10); the first body weight measurement was recorded at 15 weeks of age. Another group of animals (n=10) born and raised at the TNZ was also investigated. Body weight was assessed from 9 to 20 weeks of age.

A two-way ANOVA with repeated measures on both factors (group × time) was used for the statistical analysis of T-head, T-tail, and HLI over the conditions investigated between the two genotypes (^+/+^ vs. ^−/−^). A two-way ANOVA was employed for the statistical analysis of body weight (group × time).

#### Real-time quantitative PCR

Thirty PWScr^m+/p−^ and PWScr^m+/p+^ mice were sacrificed by cervical dislocation at three different time points: at ZT 6 (T0), immediately after 6-h of total SD (T1), and 1 h after previous SD (ZT 7; T2). Total RNA was extracted from the hypothalamus by the Trizol method (Sigma-Aldrich) according to the manufacturer’s instructions [72]. RNA concentrations were then determined by a NanoDrop 2000c spectrophotometer. The complementary DNA was obtained from up to 2 mg of total RNA by using a high-capacity RNA-to-cDNA kit (Invitrogen) and then analysed with SYBR GREEN qPCR mix. Reactions were performed in three technical replicates using an AB 7900HT fast real-time PCR system (Applied Biosystems). The relative expression levels were quantified according to the previously described ^ΔΔ^CT calculation method [64]. *Gapdh* was used as a reference gene. The specific primer pairs used for the analysis were designed using Primer3 (Supplementary Tables 5 and 6). An unpaired t-test was used to compare the differences between the genes investigated in the two genotypes.

Narcoleptic mice were sacrificed between ZT 6 and ZT 7, and the hypothalamus and cortex of the brain were dissected and immediately frozen.

#### Perfusion and immunohistochemistry (IHC)

PWScr^m+/p−^ and PWScr^m+/p+^ mice were transcardially perfused with 10 ml of PBS before infusion of 4% PFA in PBS. Perfused brains were postfixed for 24 h in 4% PFA at 4°C before immersion in 30% sucrose. To quantify lateral hypothalamic neurons expressing OX and MCH neurons, serial cryosections were cut coronally at 40 μm intervals to include brain regions within–1.10/–1.90 mm of the bregma. Before IHC staining, the sections were washed three times with PBS for 10 min each and then blocked in 5% of the appropriate serum (normal goat serum or normal donkey serum) in 0.1% Triton X-100 in PBS for 1 h. The sections were incubated with rabbit PPOX polyclonal (1:20; Millipore) and pro-melanin-concentrating hormone (PMCH; 1:50; Invitrogen) primary antibodies in 1% serum in 1% Triton X-100/PBS at room temperature overnight. The sections were then washed three times in PBS for 5 min each. They were incubated with appropriate secondary antibodies (1:1000) (goat anti-rabbit IgG, Alexa Fluor 488) in 1% serum in 1% Triton X-100/PBS for 2 h at room temperature in the dark, washed three times in PBS for 5 min each, and counterstained with Hoechst (1:400 in PBS; Sigma-Aldrich). Finally, the sections were washed in PBS and mounted on glass slides using ProLong gold antifade reagent (Invitrogen). The sections were imaged with an upright Widefield Epifluorescence Olympus BX51 microscope equipped with a 4x UPLFLN N.A objective. The microscope was controlled by Neurolucida software.

The percentage of OX- and MCH-positive cells was manually scored using NIH ImageJ software. A total of 3-5 sections were evaluated for each mouse (n= 4 for genotype), and the neuron counts were normalized to the total number of DAPI-stained nuclei (approximately 300 nuclei per microscopic field). All digital images were processed in the same way between experimental conditions to avoid artificial manipulation between different data sets. An unpaired t-test was used to compare differences between the two genotypes.

#### Chromatin immunoprecipitation (ChIP)

For the analysis of PEG3 binding, H3K4me2/3 and H3K27me3, ChIP was performed on formaldehyde cross-linked chromatin isolated from the hypothalamus of 28 PWScr^m+/p−^ and PWScr^m+/p+^ mice. Briefly, the tissue of seven different hypothalamic was minced. Formaldehyde was added to the tissue, which was resuspended in PBS at a final concentration of 1% and incubated at room temperature for 15 min while shaking. The reaction was stopped by the addition of glycine to a final concentration of 0.125 M. The tissue was washed twice with ice-cold PBS, centrifuged and resuspended in lysis buffer 1 (50 mM HEPES pH 8, 10 mM NaCl, 1 mM EDTA, 10% glycerol, 0.5% NP-40 and 0.25% Triton X-100) for 90 min at 4°C. Isolated nuclei were lysed in lysis buffer 2 (10 mM Tris–HCl pH 8.0, 200 mM NaCl, 1 mM EDTA and 0.5 mM EGTA) for 60 min at 4°C. The chromatin was sheared in sonication buffer (10 mM Tris–HCl pH 8.0, 100 mM NaCl, 1 mM EDTA, 0.5 mM EGTA, 0.1% sodium deoxycholate and 0.5% N-lauroylsarcosine) to an average size of 100–400 bp using the Sonifier 150 (Branson).

For each IP, 100 μg of sonicated chromatin was diluted in a final volume of 600 μl with sonication buffer and pre-cleared with 30 μl protein A/G agarose beads (Santa Cruz) for 4 h at 4°C on a rotating wheel. Anti-PEG3 antibody (7 μg, Abcam Ab99252), anti-histone H3K4me3 (7 μg, Abcam Ab8895), and anti_Histone H3K27me3 (7 μg, Abcam Ab195477) or rabbit IgG were added to the pre-cleared chromatin and incubated overnight at 4°C on a rotating wheel. Chromatin was precipitated with 30 μl protein A/G agarose beads for 4 h at 4°C with rotation. The beads were then washed five times with 500 μl RIPA buffer (10 mM Tris–HCl pH 8.0, 140 mM NaCl, 1 mM EDTA, 0.5 mM EGTA, 0.1% sodium deoxycholate, 1% Triton X-100 and 0.1% SDS) and once with each of the following buffers: WASH buffer (50 mM HEPES, 0.5% sodium deoxycholate, 1% Triton X-100, 1 mM EDTA, 500 mM NaCl and 0.2% NaN3), LiCl buffer (0.25 M LiCl, 0.5% NP-40, 0.5% sodium deoxycholate, 1 mM EDTA and 10 mM Tris pH 8) and TE buffer (10 mM Tris pH 8, 1 mM EDTA). The bound chromatin was eluted in 100 μl TE buffer. Crosslinks were reversed by incubation with O/N at 65°C after the addition of 1 μl RNAse cocktail (Ambion) and 2 h at 50°C after the addition of 2.5 μl SDS 20% + 2.5 μl 20 mg/ml proteinase K (Sigma). DNA was extracted by using a QIAquick Gel Extraction Kit (Qiagen). Immunoprecipitated or 1% input DNAs were analysed by real-time PCR using SYBRGreen PCR Master Mix (Bio-Rad) on a 7900HT Fast Real-Time PCR System (Applied Biosystems). Each reaction was performed in triplicate, and experiments were performed twice. The specific primer pairs used for the analysis were designed using Primer3 (Supplementary Table 7). An unpaired t-test was used to compare differences between the two genotypes.

#### Hypothalamic cell culture transfected with *Snord116*-siRNA

Embryonic rat hypothalamus cell line R7 (rHypoE-7 – Tebu-bio) cells were cultured in Dulbecco’s modified Eagle’s medium (Sigma Co. Ltd, St Louis, MO, USA) supplemented with 10% foetal bovine serum, 50 units of penicillin and 50 μg/ml of streptomycin at 37°C under an atmosphere of 5% CO2. Cells were seeded in six-well plates (Nunc Co., Roskilde, Denmark) at a density of 1 × 10^5^ cells per well. The day after cell seeding, 20 μM of Silencer® Snord116-siRNA (Ambion®) was mixed with RNAiMAX Lipofectamine (Life Technologies) reagent, and the mixture was added to each dish. We used MISSION® siRNA universal negative control #1 (Sigma) as a negative control at 20 μM. Cells were incubated for 48-h after transfection, and RNA was extracted by using the Trizol method for gene expression analysis (Supplementary Table 6). Experiments were conducted in triplicate. An unpaired t-test was used to compare differences between the two genotypes.

### RNA sequencing and analysis

Total RNA was homogenized in Trizol Reagent (Sigma-Aldrich). Libraries were prepared using a TruSeq polyA mRNA kit (Illumina) according to the manufacturer’s instructions and sequenced by using the NovaSeq 6000 System (Illumina). Raw sequence reads were quality controlled through FASTQC (https://www.bioinformatics.babraham.ac.uk/projects/fastqc/) and trimmed using Trimmomatic (v0.38) (http://www.ncbi.nlm.nih.gov/pubmed/24695404). To quantify the transcript abundances, we used Kallisto (v0.44.0) [73]. Kallisto was also used to build an index from the mouse reference genome (EnsDb.Mmusculus.v79) with default parameters. To import and summarize transcript-level abundance of Kallisto, the R package Tximport was used. Differential expression was assessed using DSeq2 [74]. Adjusted p-values less than 0.05 were selected, and only genes with > 2-fold change were considered in the analyses.

GO analysis was performed using Metascape [75] (http://metascape.org/). The identified DEGs are listed in Tables S8 and S9.

### Electrophysiological data analysis

Cortical EEG/EMG signals were recorded using Dataquest A.R.T. (Data Science International). Signals were digitized at a sampling rate of 500 Hz with a filter cut-off of 50 Hz. EEG signals were filtered at 0.3 Hz (low-pass filter) and 0.1 KHz (high-pass filter). The polysomnographic recordings were visually scored offline using SleepSign software (Kissei Comtec Co. Ltd, Japan) per four second epoch window to identify wakefulness (W), NREM or REM sleep stages, as previously described [64, 65]. Scoring was performed by a single observer who was blinded to the mouse groups.

Specifically, W, NREM and REM states were scored when characteristic EEG/EMG activity occupied 75% of the epoch [64, 65]. EEG epochs determined to have artefact (interference caused by scratching, movement, eating, or drinking) were excluded from the analysis. Artefact comprised <5-8% of all recordings used for analysis.

The percentage of time spent in total sleep, NREM and REM sleep out of the total recording time was determined. The amount of time spent in each stage was established by counting the types of epochs (W or NR or R) and averaging over 2-h periods. The spectral characteristics of the EEG were further analysed. The EEG power densities of the delta (0, 5–4 Hz) and theta (5–9 Hz) frequencies in NREM and REM sleep were computed for all conditions investigated. To exclude variability due to the implantation, the power density of each animal was normalized to the power density of the last 4-h of the light period of the 24-h recording (22°C and 30°C, respectively). The temperature was recorded at a sampling rate of 5 Hz within the range of 34°C – 41°C and averaged over 2-h periods. A two-way ANOVA with repeated measures (factors of group × time) was used for the statistical analysis of the 2-h averaged time-course changes in the percentage of each sleep stage (W, NREM and REM sleep), delta power, theta power and body temperature between the two genotypes (^+/+^ vs. ^−/−^). The statistical analysis of the cumulative amount of W, NREM, and REM sleep over the dark and light periods among the groups at BL, after SD and at the TNZ was performed with one-way ANOVA. An unpaired t-test was used to compare differences in the sleep stages and in the delta and theta power during the first 4 h of rebound after SD between the two genotypes.

SUA data acquisition was performed using an RX7 system (Tucker Davis Technology, TDT). The synchronization between the behavioural set up and the TDT system was guaranteed with transistor-transistor logic (TTL) from the CHORA feeders to the TDT system. The data were digitally sampled at 12 kHz. To detect spike timestamps, the neural traces were filtered with a bandpass filter (300 – 5000 Hz), and then, the common average reference (CAR) was applied [66]. Spikes were detected using a hard threshold that was computed as previously described [67]. A correlation filter was applied between each detected spike and the corresponding signal chunk in the other recording sites to detect and exclude false positive spikes given by movement artefact.

To specifically identify active neurons in the LH, we imposed two criteria: (i) the refractory period following a spike was set to 1 ms; and (ii) the maximum duration of a waveform was set to 5 ms. Spikes of individual neurons were sorted offline using the noise robust LDA-GMM algorithm based on a linear discriminant analysis [68]. To identify neurons that specifically respond to a particular sleep-wake cycle, the unit activity was subsequently analysed per 4-s epoch in each sleep-wake stage for the average discharge rate (spikes per second). Classification of units according to the state in which their maximal discharge rate occurred was performed by ANOVA followed by post hoc paired t-tests with the Bonferroni correction, (P <0.05)[69]. This analysis allowed the classification of units into three main neuronal populations that were shown to respond during specific sleep-wake stages (Figure 1B).

Moreover, to identify the neurons related to feeding behaviour in the LH, we assessed the firing rate of neurons that responded after the food was released by aligning the unit firing to the pellet release from the CHORA feeder after a spontaneous nose-poke activity. The mean baseline firing rate of each neuron was determined in the interval of 5 s before the pellet was released. Paired Student’s t-tests were used to classify the firing rate differences of the same units before and after the food was released. To motivate the mice to perform more trials, they were food deprived for 12 h during the dark period, and the recording session started at the beginning of the light period for the following 2 h. Sessions were videotaped and reviewed to eliminate trials in which the mice performed the nose-poke activity but the food was not eaten. Video analysis also demonstrated that grooming rarely occurred and, thus, did not affect SUA. Indeed, grooming has been shown to induce moderate OX neuron activity [70]. Among all sessions, a mean of 3.25 ± 1.49 trials for the PWScr^m+/p−^ mice and 2 ± 0.69 trials for the PWScr^m+/p−^ mice were removed from the analysis because the food was not eaten.

### Statistics

Values were tested for a Gaussian distribution with the Kolmogorov–Smirnov test. Data are presented as the mean ± standard error of the mean (SEM). Two-way repeated-measures analysis of variance (ANOVA) was used to perform group comparisons with multiple measurements. Paired and unpaired t tests were used for single value comparisons. One-way ANOVA was used to compare more than two groups, followed by post hoc Tukey’s test. The Bonferroni correction was further applied in the post hoc analysis, as appropriate, to correct for multiple comparisons. Neuronal dynamics of the LH were analysed by the chi-square test. Phenopy [61] was used for sleep, temperature and SUA analyses, while GraphPad Prism6 (GraphPad Prism Software, Inc.) was used for statistical analysis. Type I error α was set to 0.05 (p < 0.05).

## Supporting information

Pace-Falappa et al. Figure Supplementary 1

Pace-Falappa et al. Figure Supplementary 2

Pace-Falappa et al. Figure Supplementary 3

Pace-Falappa et al. Figure Supplementary 4

Pace-Falappa et al. Figure Supplementary 5

Pace-Falappa et al. Figure Supplementary 6

Pace-Falappa et al. Table S1

Pace-Falappa et al. Table S2

Pace-Falappa et al. Table S3

Pace-Falappa et al. Table S4

Pace-Falappa et al. Table S5

Pace-Falappa et al. Table S6

Pace-Falappa et al. Table S7

Pace-Falappa et al. Table S8

Pace-Falappa et al. Table S9

Pace-Falappa et al. Supplementary materials

## Author contributions

MP and VT designed the study. MP, MF performed the animal experiments. VT provided infrastructural support. AF and MP performed the gene expression analysis. MP and MF performed the EEG and SUA analysis. AF and MP analysed the gene expression data. AF performed the ChiP experiment. MC analysed data from the infrared thermocamera. AU FK and EK analysed the RNAseq. CB,and ZG provide and dissected brain samples from narcoleptic mice. MP, MF, MC, RA, AU and VT drafted and finalized the manuscript. All authors revised and finalized it.

## Acknowledgement

We thank Simone Bellini, a master student of Tucci’s laboratory for her help with the RNA extraction and RT-qPCR. This study was financially supported by Jerome Lejeune Foundation under Grant Agreements No 1599-TV2016B, by the The Foundation for Prader Willi Research. Additionally, this project received the Seal of Excellence by the European Union’s Horizon 2020, N 753417.

