## Supplementary figures and images for "Paternally expressed imprinted *Snord116* and *Peg3* regulate hypothalamic orexin neurons"

### Pace-Falappa et al. Figure Supplementary 1

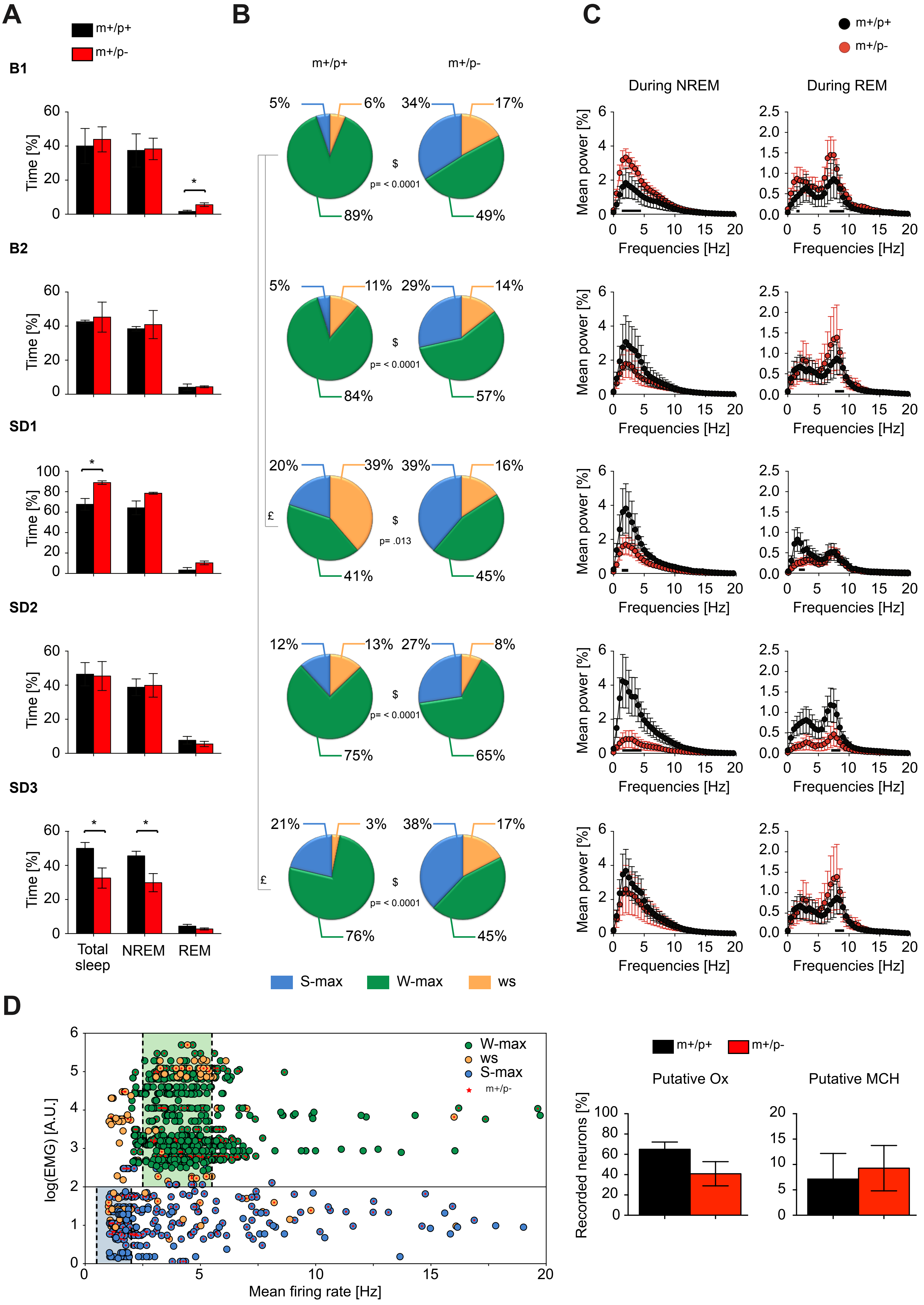

### Pace-Falappa et al. Figure Supplementary 2

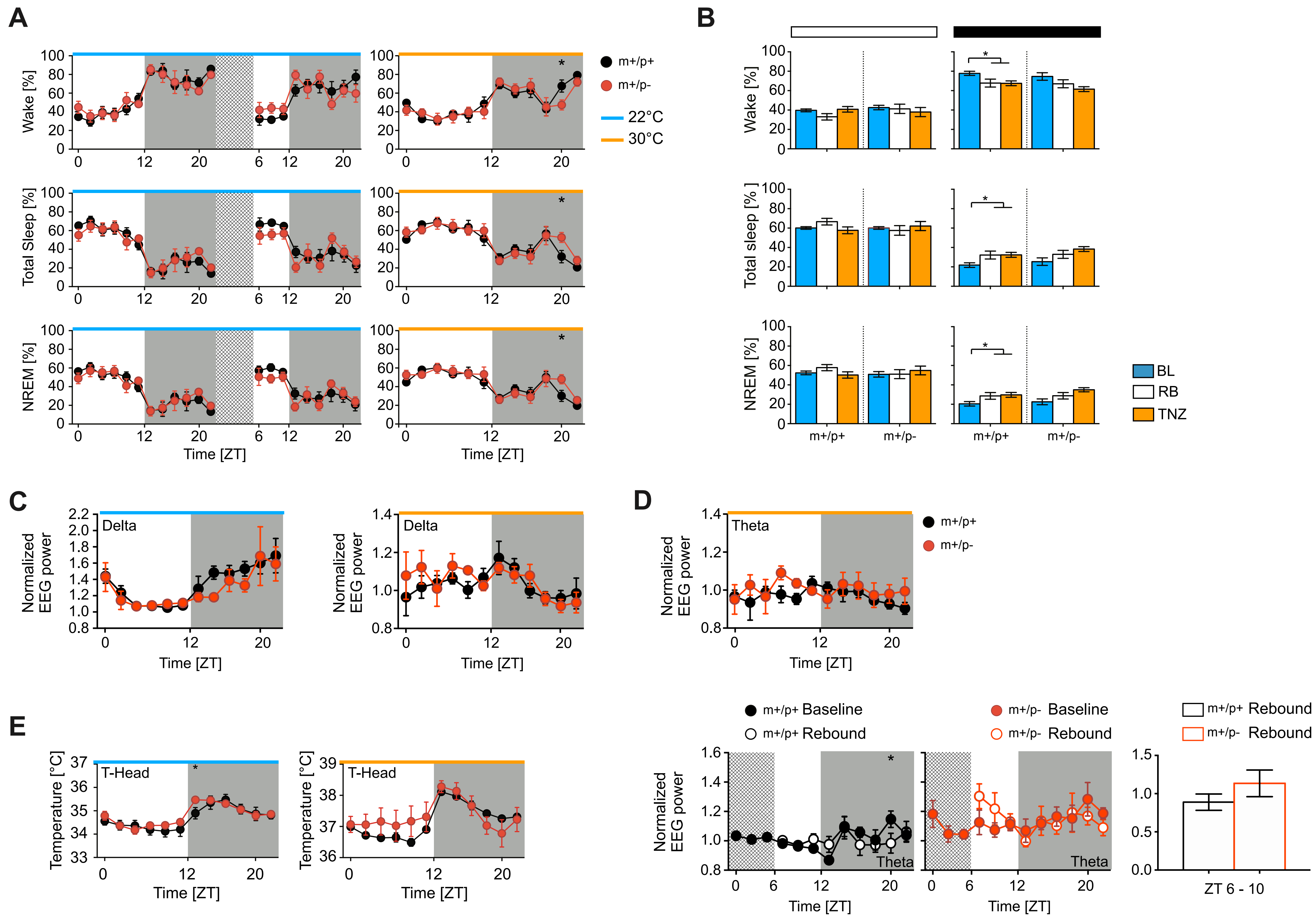

### Pace-Falappa et al. Figure Supplementary 3

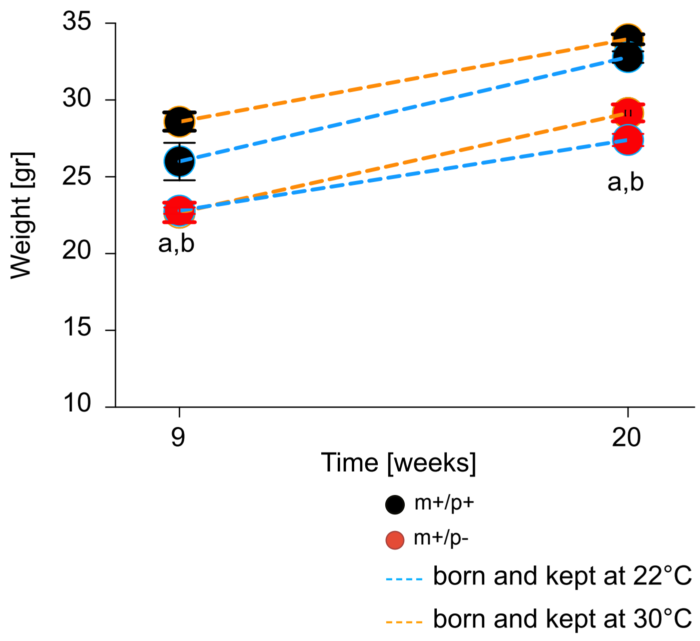

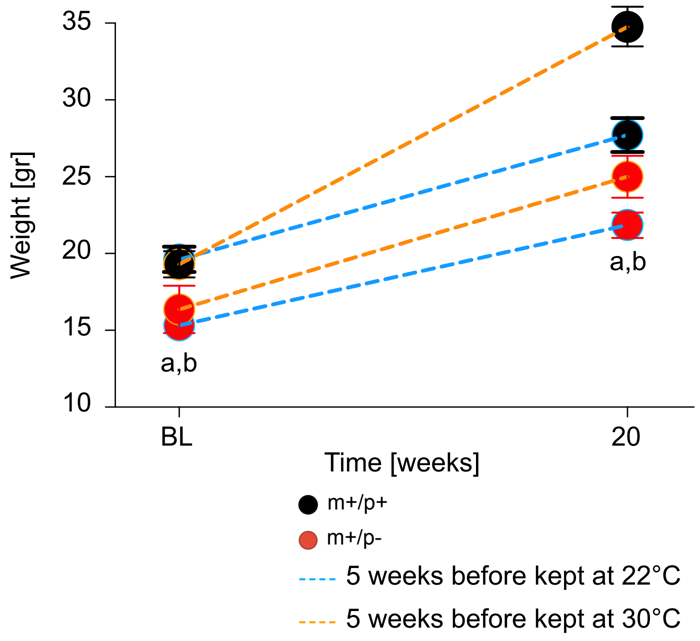

### Pace-Falappa et al. Figure Supplementary 4

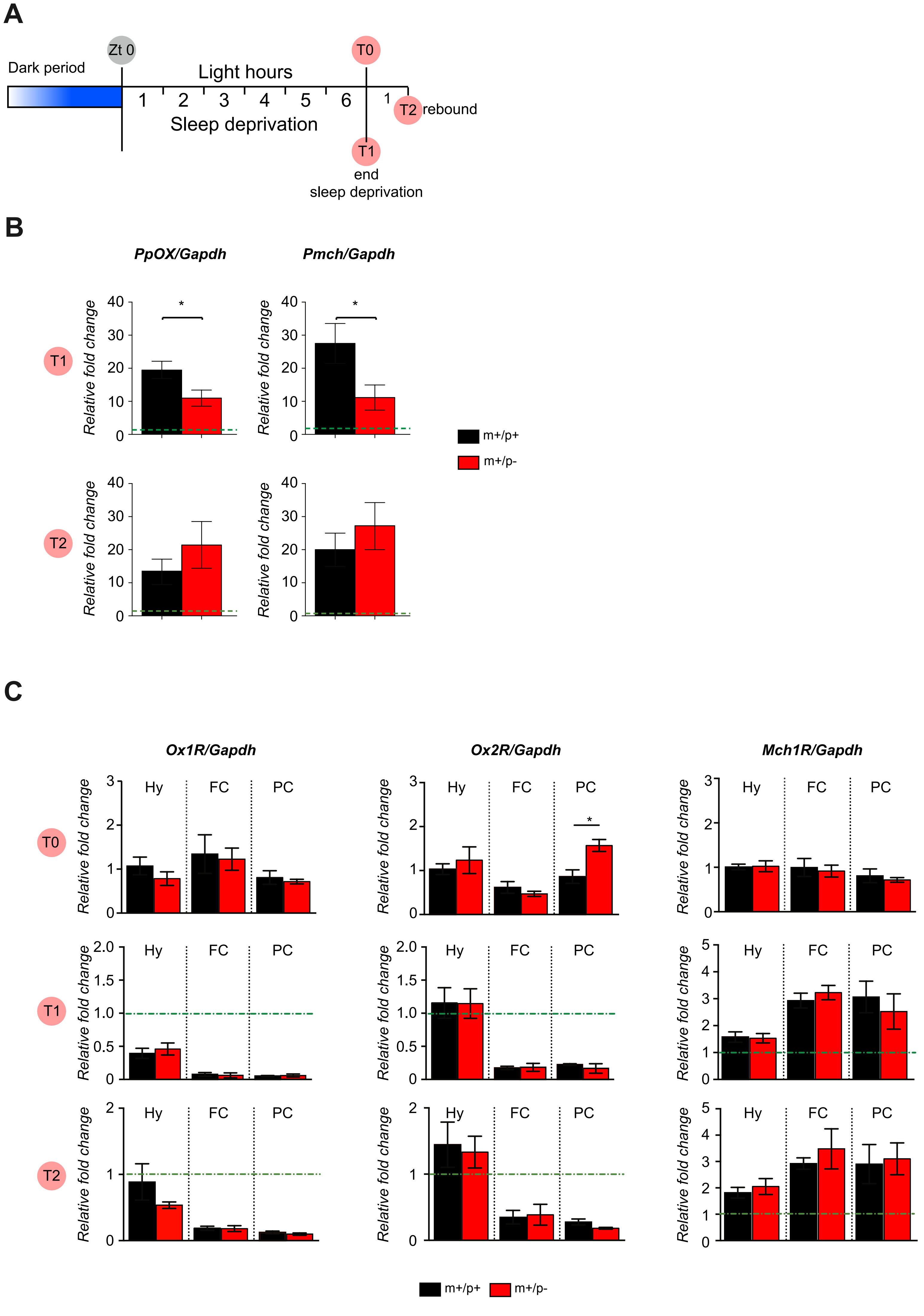

### Pace-Falappa et al. Figure Supplementary 5

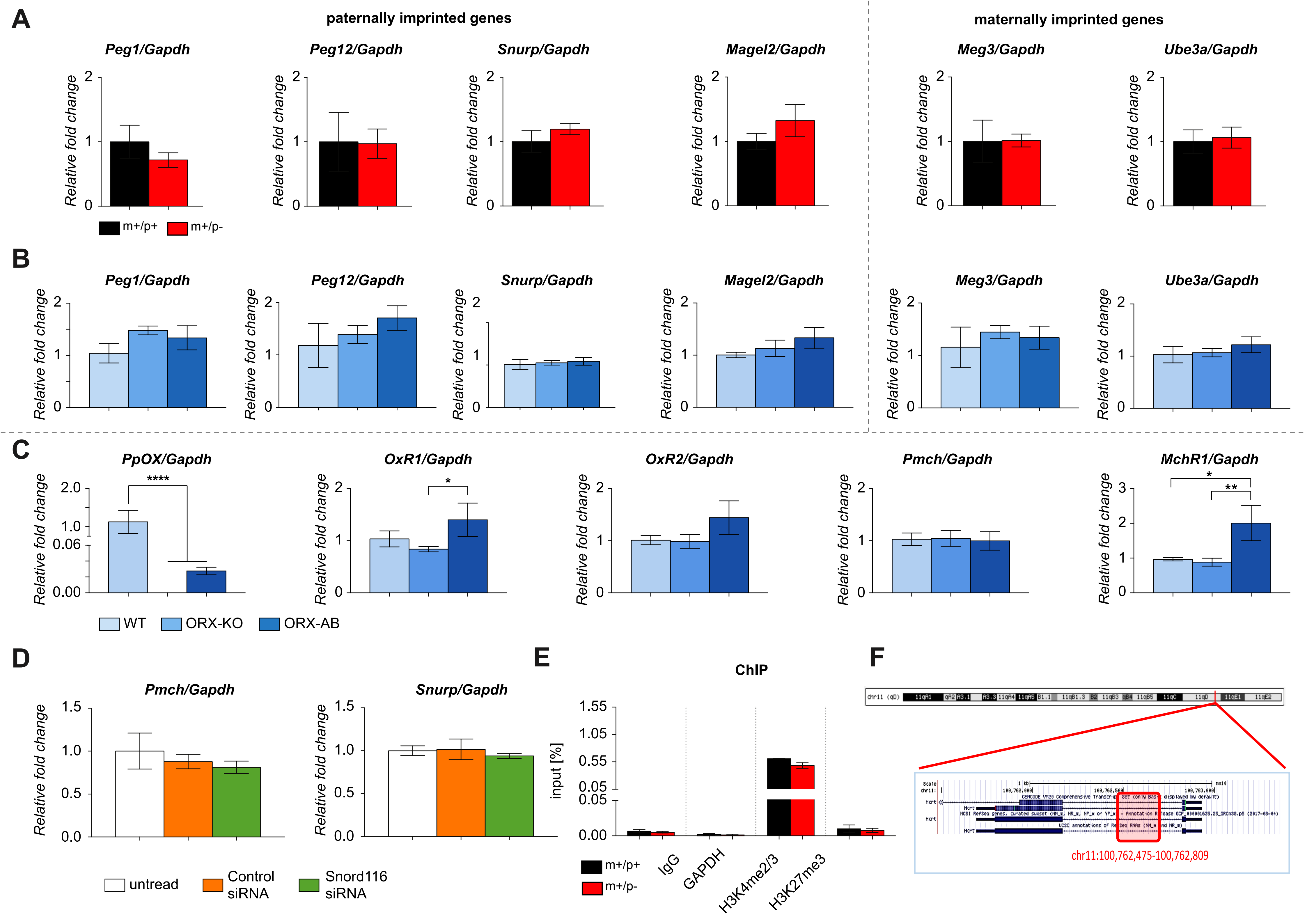

### Pace-Falappa et al. Figure Supplementary 6

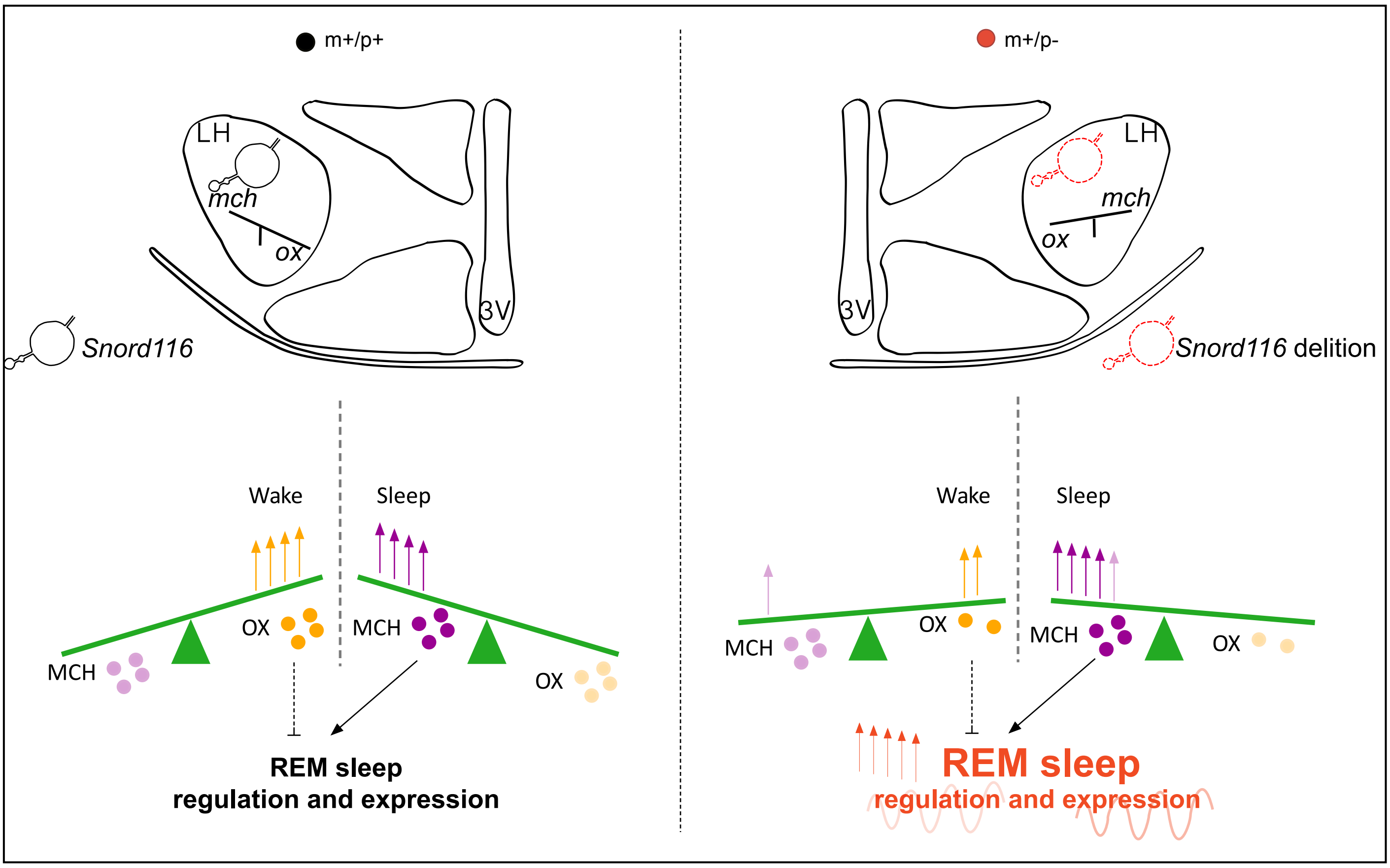
