## Supplementary material for "Paternally expressed imprinted *Snord116* and *Peg3* regulate hypothalamic orexin neurons": Pace-Falappa et al. Table S1

| Phase: | B1 | | B2 | | SD | | SD1 | | SD2 | |
|---|---|---|---|---|---|---|---|---|---|---|
| Genotype: | m+/p+ | m+/p- | m+/p+ | m+/p- | m+/p+ | m+/p- | m+/p+ | m+/p- | m+/p+ | m+/p- |
| W-max | 90 | 60 | 104 | 43 | 48 | 55 | 77 | 73 | 96 | 54 |
| **Wake** | | | | | | | | | | |
| NR - max | 2 | 21 | 1 | 19 | 6 | 27 | 6 | 17 | 24 | 23 |
| R - max | 2 | 4 | 2 | 4 | 2 | 17 | 5 | 5 | 2 | 11 |
| NRR - max | 1 | 5 | 3 | 7 | 15 | 3 | 1 | 9 | 1 | 12 |
| **Sleep** | | | | | | | | | | |
| wsp | 9 | 11 | 1 | 9 | 14 | 11 | 13 | 5 | 4 | 16 |
| WR - max | 3 | 4 | 6 | 6 | 31 | 8 | 0 | 4 | 0 | 5 |
| **None** | | | | | | | | | | |
| Total | 107 | 105 | 117 | 88 | 116 | 121 | 102 | 113 | 127 | 121 |

| Genotype: | m+/p+ | m+/p- |
|---|---|---|
| W-max | 415 | 285 |
| **Wake** | | |
| NR - max | 39 | 107 |
| R - max | 13 | 41 |
| NRR - max | 21 | 36 |
| **Sleep** | | |
| wsp | 41 | 52 |
| WR - max | 40 | 27 |
| **None** | | |
| Total | 569 | 548 |
