## Supplementary material for "Paternally expressed imprinted *Snord116* and *Peg3* regulate hypothalamic orexin neurons": Pace-Falappa et al. Table S2

| B1 | m+/p+ | m+/p- | Totals |
|---|---|---|---|
| W-Max | 90 | 60 | 150 |
| Others | 17 | 45 | 62 |
| Totals | 107 | 105 | 212 |
| | | p-value | <.0001 |

| | m+/p+ | m+/p- | Totals |
|---|---|---|---|
| S-max | 5 | 30 | 35 |
| Others | 102 | 75 | 177 |
| Totals | 107 | 105 | 212 |
| | | p-value | <.0001 |

| | m+/p+ | m+/p- | Totals |
|---|---|---|---|
| ws | 12 | 15 | 27 |
| Others | 95 | 90 | 185 |
| Totals | 107 | 105 | 212 |
| | | p-value | ns |

| B2 | m+/p+ | m+/p- | Totals |
|---|---|---|---|
| W-Max | 104 | 43 | 147 |
| Others | 13 | 45 | 58 |
| Totals | 117 | 88 | 205 |
| | | p-value | <.0001 |

| | m+/p+ | m+/p- | Totals |
|---|---|---|---|
| S-max | 6 | 30 | 36 |
| Others | 111 | 58 | 169 |
| Totals | 117 | 88 | 205 |
| | | p-value | <.0001 |

| | m+/p+ | m+/p- | Totals |
|---|---|---|---|
| ws | 7 | 15 | 22 |
| Others | 110 | 73 | 183 |
| Totals | 117 | 88 | 205 |
| | | p-value | .02 |

| SD | m+/p+ | m+/p- | Totals |
|---|---|---|---|
| W-Max | 48 | 55 | 103 |
| Others | 68 | 66 | 134 |
| Totals | 116 | 121 | 237 |
| | | p-value | ns |

| | m+/p+ | m+/p- | Totals |
|---|---|---|---|
| S-max | 23 | 47 | 70 |
| Others | 93 | 74 | 167 |
| Totals | 116 | 121 | 237 |
| | | p-value | .0017 |

| | m+/p+ | m+/p- | Totals |
|---|---|---|---|
| ws | 45 | 19 | 64 |
| Others | 71 | 102 | 173 |
| Totals | 116 | 121 | 237 |
| | | p-value | <.0001 |

| SD1 | m+/p+ | m+/p- | Totals |
|---|---|---|---|
| W-Max | 77 | 73 | 150 |
| Others | 25 | 40 | 65 |
| Totals | 102 | 113 | 215 |
| | | p-value | ns |

| | m+/p+ | m+/p- | Totals |
|---|---|---|---|
| S-max | 12 | 31 | 43 |
| Others | 90 | 82 | 172 |
| Totals | 102 | 113 | 215 |
| | | p-value | .059 |

| | m+/p+ | m+/p- | Totals |
|---|---|---|---|
| ws | 13 | 9 | 22 |
| Others | 89 | 104 | 193 |
| Totals | 102 | 113 | 215 |
| | | p-value | ns |

| SD2 | m+/p+ | m+/p- | Totals |
|---|---|---|---|
| W-Max | 96 | 54 | 150 |
| Others | 31 | 67 | 98 |
| Totals | 127 | 121 | 248 |
| | | p-value | <.0001 |

| | m+/p+ | m+/p- | Totals |
|---|---|---|---|
| S-max | 27 | 46 | 73 |
| Others | 100 | 75 | 175 |
| Totals | 127 | 121 | 248 |
| | | p-value | .052 |

| | m+/p+ | m+/p- | Totals |
|---|---|---|---|
| ws | 4 | 21 | 25 |
| Others | 123 | 100 | 223 |
| Totals | 127 | 121 | 248 |
| | | p-value | .0002 |
