## Supplementary material for "Paternally expressed imprinted *Snord116* and *Peg3* regulate hypothalamic orexin neurons": Pace-Falappa et al. Table S3

| m+/p+ | **B1** | **B2** | Totals |
|---|---|---|---|
| ws | 12 | 7 | 19 |
| W-max | 90 | 104 | 194 |
| S-max | 5 | 6 | 11 |
| Totals | 107 | 117 | 224 |
| | | p-value | ns |

| m+/p- | **B1** | **B2** | Totals |
|---|---|---|---|
| ws | 15 | 15 | 30 |
| W-max | 60 | 43 | 103 |
| S-max | 30 | 30 | 60 |
| Totals | 105 | 88 | 193 |
| | | p-value | ns |

| m+/p+ | **B1** | **SD** | Totals |
|---|---|---|---|
| ws | 12 | 45 | 57 |
| W-max | 90 | 48 | 138 |
| S-max | 5 | 23 | 28 |
| Totals | 107 | 116 | 223 |
| | | p-value | < .0001 |

| m+/p- | **B1** | **SD** | Totals |
|---|---|---|---|
| ws | 15 | 19 | 34 |
| W-max | 60 | 55 | 115 |
| S-max | 30 | 47 | 77 |
| Totals | 105 | 121 | 226 |
| | | p-value | ns |

| m+/p+ | **B1** | **SD1** | Totals |
|---|---|---|---|
| ws | 12 | 13 | 25 |
| W-max | 90 | 77 | 167 |
| S-max | 5 | 12 | 17 |
| Totals | 107 | 102 | 209 |
| | | p-value | ns |

| m+/p- | **B1** | **SD1** | Totals |
|---|---|---|---|
| ws | 15 | 9 | 24 |
| W-max | 60 | 74 | 134 |
| S-max | 30 | 31 | 61 |
| Totals | 105 | 114 | 219 |
| | | p-value | ns |

| m+/p+ | **SD1** | **SD2** | Totals |
|---|---|---|---|
| ws | 12 | 4 | 16 |
| W-max | 90 | 96 | 186 |
| S-max | 5 | 27 | 32 |
| Totals | 107 | 127 | 234 |
| | | p-value | .0001 |

| m+/p- | **B1** | **SD2** | Totals |
|---|---|---|---|
| ws | 15 | 21 | 36 |
| W-max | 60 | 54 | 114 |
| S-max | 30 | 46 | 76 |
| Totals | 105 | 121 | 226 |
| | | p-value | ns |

Table S3
