## Supplementary material for "Paternally expressed imprinted *Snord116* and *Peg3* regulate hypothalamic orexin neurons": Pace-Falappa et al. Table S4

| | m+/p+ | m+/p- | Totals |
|---|---|---|---|
| Type I | 28 | 16 | 44 |
| Others | 119 | 58 | 177 |
| Totals | 147 | 74 | 221 |
| | | p-value | ns |

| | m+/p+ | m+/p- | Totals |
|---|---|---|---|
| Type II | 105 | 22 | 127 |
| Others | 42 | 52 | 94 |
| Totals | 147 | 74 | 221 |
| | | p-value | < .0001 |

| | m+/p+ | m+/p- | Totals |
|---|---|---|---|
| Type III | 14 | 36 | 50 |
| Others | 133 | 38 | 171 |
| Totals | 147 | 74 | 221 |
| | | p-value | < .0001 |
