## Supplementary material for "Paternally expressed imprinted *Snord116* and *Peg3* regulate hypothalamic orexin neurons": Pace-Falappa et al. Table S5

### OREXIN SYSTEM

| GENE | FORWARD | REVERSE |
|---|---|---|
| PPHCRT | 5'- CAGGCACCATGAACTTTCCTTCTA | 5'- CAGCAGCAGCGTCACG |
| OREXIN R1 | 5'- CGCCAACCCTATCATCTACAA | 5'- GCTCTGCAAGGACAAGGACTT |
| OREXIN R2 | 5'- GCTCACCAGCATAAGCACACT | 5'- TGGACAGGAGTGAAGATGGTACT |

### MCH SYSTEM

| GENE | FORWARD | REVERSE |
|---|---|---|
| PMCH | 5'- ATTCAAAGAACACAGGCTCCAAAC | 5'- CGGATCCTTTCAGAGCAAGGTA |
| MCH R1 | 5'- GCCACCTCCTCGCACAA | 5'- CTTCACCACGGCAAAAATGAC |

### MATERNAL IMPRINTED GENES

| GENE | FORWARD | REVERSE |
|---|---|---|
| UBE3A | 5'- GCGAGCAGCTGCAAAGCATCTAAT | 5'- AGCTTGCTCCTTTCTTGGAGGGAT |
| MEG3 | 5'- CACAGAAGACGAAGAGCTGGA | 5’- GGTAGAGGTGCACAGCAGGT |

### PATERNAL IMPRINTED GENES

| GENE | FORWARD | REVERSE |
|---|---|---|
| PEG1 | 5’- CCATATTTGAGCAGGCCAGC | 5’- TCCATTCGAACAGACAGAGACT |
| PEG3 | 5’- GACTCGTCCTCACGATCCC | 5’- AGTCCAGCTTGCCGAAGAT |
| PEG12 | 5’- GGCACAGCTCAGAACTAGGG | 5’- AAGTCCTGGGTGAATCCCTT |
| MAGEL2 | 5’- AGATTTTTGGCCGCACTCTA | 5’- GGCATTTGCGTCCTTGTATT |
| SNRPN/SNURF | 5’- CTGCTCTTCCAACCCCAGG | 5’- ATGCAAAACAGCCAGAACGT |

### HOUSEKEEPING GENE

| GENE | FORWARD | REVERSE |
|---|---|---|
| GAPDH | 5'- GAACATCATCCCTGCATCCA | 5'- CCAGTGAGCTTCCCGTTCA |
