## Supplementary material for "Paternally expressed imprinted *Snord116* and *Peg3* regulate hypothalamic orexin neurons": Pace-Falappa et al. Table S6

### CHIP Primers

| GENE | FORWARD | REVERSE |
|---|---|---|
| PPHCRT | 5’ – TCCTTTGCTGGGGAAACTGT – 3 | 5’ – AGAGAGGAGTATGGGTGGGT – 3’ |
| SNRPN | 5’ – TAACACACCCAAGGAGTCCG – 3 | 5’ – GACTAGCGCAGAGAGAGGAGAG – |
| GAPDH | 5’ – CCCAGCCAAGTTTGAAAGGG – 3 | 5’ – GCATCTCCCTCACACACCTCTT – 3’ |
