## Supplementary material for "Paternally expressed imprinted *Snord116* and *Peg3* regulate hypothalamic orexin neurons": Pace-Falappa et al. Table S7

### RATS primers

| GENE | FORWARD | REVERSE |
|---|---|---|
| PPHCRT | 5’ – AACCTTCCTTCTACAAAGGTTCC – 3’ | 5’ – CAGCTCCGTGCAACAGTTC– 3’ |
| PMCH | 5’ – CACAAAGAACACAGGCTCCA – 3’ | 5’ – TTCCCTCTTTTCCTGTGTGG – 3’ |
| Peg3 | 5’ – GGGGAGTGCTACCTTCTTGA – 3’ | 5’ – CTGTTTTGCTCACACCCAAG – 3’ |
| Snord116 | 5’ – TGCTTGGATCGATGATGATTT – 3’ | 5’ – CTGGACCTCAGTCACGATGAT – 3’ |
| Snrpn/Snurf | 5’ – CAGCAATCATGACTGTGGGTA – 3’ | 5’ – TCTTTGGCTTGATCTTCCTGA – 3’ |
| Gapdh | 5’ – GAACATCATCCCTGCATCCA – 3’ | 5’ – CCAGTGAGCTTCCCGTTCA – 3’ |
