## Supplementary material for "Paternally expressed imprinted *Snord116* and *Peg3* regulate hypothalamic orexin neurons": Pace-Falappa et al. Supplementary materials

**General information:**

Abstract words: 186; words count: 9525; figures: 4; Supplementary material: figures 6; table 8

**Conflict of Interest: each author discloses the absence of any conflicts of interest relative to the research covered in the submitted manuscript.**

**Figure S1**

**PWScr^m+/p−^ mice have altered neuronal dynamics of the LH in relation to sleep and food**. The results of single unit activity (SUA) combined with EEM/EMG recordings. **A)** The percentage of total time spent in sleep stages (NREM plus REM sleep), NREM sleep and REM sleep, in PWScr^m+/p−^ mice (red bar) compared with control mice (black bar). EEG/EMG/SUA were recorded at different time points and are shown in each row: B1 and B2, baseline; SD1, 1-h immediately after 6 h of SD; and SD2 and SD3, two time points during the 18-h recovery period. At B1, REM sleep was increased in PWScr^m+/p−^ mice (unpaired t-test: t(6) = 3.06, p = .02) relative to PWScr^m+/p+^ mice. At B2, no differences between the two genotypes were observed in any of the sleep stages investigated. At SD1, total sleep was increased in PWScr^m+/p−^ mice (unpaired t-test: t(6) = 3.53, p = .01) relative to controls. At SD2, no differences between the two genotypes were observed in any of the sleep stages investigated. At SD3, total sleep (unpaired t-test: t(6) = 2.52, p = .04) and NREM sleep (unpaired t-test: t(6) = 2.64, p = .03) were decreased in PWScr^m+/p−^ mice relative to PWScr^m+/p+^ mice. Data are presented as the mean ± SEM. Asterisks (*) indicate a significant difference between genotypes: * P ≤ .05. **B)** The panel shows the neuronal distribution of recorded neurons (according to the classification described in the Methods session) in a pie chart for each time point. W-max neurons are shown in green, S-max neurons are shown in blue, and ws neurons are shown in yellow. Differences between the two genotypes are indicated by $, while differences within groups across time points are indicated by §. Significance was computed with the chi-square test. **C)** The normalized power density of the whole spectrum during NREM and REM sleep (left and right, respectively) for both genotypes; red circles depict PWScr^m+/p−^ mice, and black circles depict PWScr^m+/p+^ mice. **D)** Neuronal classification in a 2D scatter plot. The x-axis plots the mean firing rate of each cell in the state in which they maximally fire, and the y-axis plots the mean logarithm of EMG signals according to the selected sleep-wake state. W-max neurons are shown in green, S-max neurons in blue, and ws neurons in yellow. Neurons recorded from mutant mice are marked by red dots. Putative OX neurons (green squares) and MCH putative neurons (blue squares) are also shown (see Methods). The two genotypes investigated were PWScr^m+/p−^ mice (n=4) and PWScr^m+/p+^ mice (n= 4).

**Figure S2**

**Sleep-wake cycle and temperature profile recorded**. **A)** The panel shows the hourly percentage of time spent in wakefulness, total sleep (including both NREM sleep and REM sleep), NREM sleep, and REM sleep in PWScr^m+/p−^ mice (red) versus control mice (black), recorded over an uninterrupted 24-h period of the 12-h light/dark cycle for baseline (BL) and during the 18-h after SD. Recordings were conducted in animals maintained at 22°C (cyan bar in each graph) and at 30°C (orange bar in each graph). No significant differences were found at 22°C between genotypes among all sleep stages investigated. At 30° C, a two-way repeated-measures ANOVA revealed significant main effects of time for wakefulness (F(11,88) = 13.70, p < .0001 “time”), for total sleep (F(11,88) = 13.51, p < .0001 “time”) and for NREM sleep (F(11,88)= 12.50, p = <.0001 “time”). Data are presented as the 2-h mean values ± SEM. **B)** Cumulative amount of time spent in wakefulness, total sleep and NREM sleep for PWScr^m+/p−^ and PWScr^m+/p+^ mice over the 12-h light period and the 12-h dark period. Cyan bar indicates the baseline (BL) value, the white bar indicates the 18-h recovery (RB) period following 6 h of SD, and the orange bar indicates recordings at 30°C. Statistical analysis was performed by one-way ANOVA followed by post hoc analysis with the Bonferroni multiple comparison test. Dark period: PWScr^m+/p+^ mice showed a difference in the percentage of wakefulness (F(1.79, 7.19)= 6.85, p = .02), total sleep (F(1.80, 7.20)= 7.35, p = .01) and NREM sleep (F(1.86, 7.46)= 5.40, p = .03). Data are presented as the mean ± SEM. Asterisks (*) indicate a significant difference between genotypes: * P ≤ .05. **C-D)** Spectral analysis. Normalized delta and theta power during NREM sleep (C-upper) and REM sleep (D-upper) at 22°C (cyan bar above the graph) and at 30°C (orange bar above the graph) and after SD and relative to baseline for PWScr^m+/p−^ mice (D-button) were recorded between genotypes. No substantial differences were observed between genotypes and conditions investigated. Data are presented as the means of 2-h bins ± SEM. Asterisks (*) indicate a significant difference between genotypes: * P ≤ .05; **p ≤ .01; *** p ≤ .001; **** p ≤ .0001. The bottom right shows the normalised theta power recorded from ZT 6 to ZT 10, and no differences were observed between genotypes. Two genotypes were investigated: PWScr^m+/p−^ mice (n= 10, 5 mice at 22°C and 5 mice at 30°C) and PWScr^m+/p+^ mice (n= 10, 5 mice at 22°C and 5 mice at 30°C). **E)** Head temperature (Head-T) profiles for PWScr^m+/p+^ and PWScr^m+/p-^ mice were recorded with an infrared thermocamera over 24 h. Recordings were performed under two different environmental conditions: at 22°C (cyan bar over the graph, shown on the left) and at 30°C, representing the thermoneutrality zone (TNZ) (orange bar over the graph, shown on the right). T-head was slightly increased in PWScr^m+/p-^ mice relative to control mice during the dark period at ZT 14 at 22°C. Asterisks (*) indicate a significant difference between genotypes: * P ≤ .05.

**Figure S3**

***Snord116* loss induces growth retardation.** Body weight was assessed at 22°C and 30°C (TNZ). **A)** The absolute weekly body weights of mice born at 22°C (cyan connection) and mice born at 30°C (orange connection) were assessed. PWScr^m+/p−^ mice are indicated by red circles, while PWScr^m+/p+^ mice are represented by black circles. Two-way ANOVA revealed that PWScr^m+/p−^ mice had significant growth retardation compared with control mice when raised at either 22°C or 30°C (F(3, 32)= 179.5, p = < .0001 “time”; F(3, 32)= 45.07, p = < .0001 “genotypes”). a indicates differences between genotypes at 22°C, while b indicates differences between genotypes at 30°C. Two genotypes were investigated: PWScr^m+/p−^ mice (n= 10, 5 mice at 22°C and 5 mice at 30°C) and PWScr^m+/p+^ mice (n= 10, 5 mice at 22°C and 5 mice at 30°C). **B)** At 15 to 20 weeks of age (at the time point when the sleep-wake cycle was recorded), the absolute weekly body weight was assessed in mice after housing for 5 weeks either at 22°C or 30°C. Two-way ANOVA revealed that PWScr^m+/p−^ mice had significant growth retardation compared with control mice housed at either 22°C or 30°C (F(3, 32)= 10.58, p = < .0001 “interaction”). a indicates differences between genotypes at 22°C, while b indicates differences between genotypes at 30°C. Two genotypes were investigated: PWScr^m+/p−^ mice (n= 10, 5 mice at 22°C and 5 mice at 30°C) and PWScr^m+/p+^ mice (n= 10, 5 mice at 22°C and 5 mice at 30°C).

**Figure S4**

**Gene expression analysis of the OX amd MCH systems**. **A)** Experimental timeline. PWScr^m+/p−^ and PWScr^m+/p+^ mice were sacrificed at three different time points to assess the gene expression of prepro-OX (*Ppox*) and the precursor of MCH (*Pmch*). Mice were sacrificed at Zeitgaber 6 (ZT 6; T0), after 6-h of SD at ZT 6 (T1), and after 6-h of SD (ZT 7; T2). **B)** At T1, PWScr^m+/p−^ mice showed reduced *Ppox* (unpaired t-test: t(8) = 3.14, p = .01) and *Pmch* (unpaired t-test: t(8) = 2.66, p = .02) compared with PWScr^m+/p+^ mice. At T2, no differences were observed between the two genotypes for both genes. **C)** Gene expression analysis of the OX receptor 1 (Ox1R), the OX receptor 2 (Ox2R) and the MCH receptor 1 (*Mch1R*) in the hypothalamus (Hy), the frontal cortex (FC) and the parietal cortex (PC). The green lines in T1 and T2 represent the baseline value (T0). Only OxR2 in the PC was found to be increased in PWScr^m+/p−^ mice (unpaired t-test: t(8) = 3.44, p = .008). Gene expression (mean ± SEM) was assessed by qRT-PCR in mice of the 2 genotypes: PWScr^m+/p−^ mice (n= 15, 5 mice for each time point) and PWScr^m+/p+^ mice (n= 15, 5 mice for each time point). Gapdh was used as a reference gene.

**Figure S5**

**Gene expression analysis of the maternally and paternally imprinted genes in PWScr^m+/p−^ mice and in mice of two narcoleptic models**. **A)** Gene expression analysis of *Peg1, Peg12, Snurp* *Magel2 Meg3* and *Ube3a* in PWScr^m+/p−^ mice relative to control mice. Unpaired t tests did not reveal any significant changes between the two genotypes. **B)** Gene expression of *Peg1, Peg12, Snurp Magel2* *Meg3* and *Ube3a* in WT, KO and Atx mice. One-way ANOVA did not reveal any significant changes between genotypes. **C)** Gene expression analysis of *Ppox, OxR1, OxR2, Pmch* and *MchR1* in WT, KO and Atx mice. We confirmed a significant reduction in *Ppox* in both strains of narcoleptic mice relative to WT mice (one-way ANOVA: F(3,16)= 11.63, Bonferroni post hoc test p = <.0001). *OxR1* (one-way ANOVA: F(3,16)= 7.034, Bonferroni post hoc test p = <.03) and *MchR1* (one-way ANOVA: F(2,15)= 5.17, Bonferroni post hoc test p = <.01) were reduced only in Atx mice relative to WT and KO mice. Gene expression (mean ± SEM) was assessed by qRT-PCR in mice of the 2 genotypes: PWScr^m+/p−^ mice (n= 5) and PWScr^m+/p+^ mice (n=5). KO (n= 12) and Atx (n=4) narcoleptic mice were investigated and compared with WT mice (n=4). *Gapdh* was used as a reference gene. **D)** Gene expression analysis of *Pmch* and *Snurp* in the *Snord116*-siRNA-treated immortalised hypothalamic rat cell line (green bars). Both genes were unchanged compared with untreated cells or scrambled siRNA-treated cells (white and orange bars). **E)** ChIP analysis of PEG3 binding on the *Gapdh* promoter region in PWScr^m+/p−^ mice (red) versus controls (black). Values are expressed as the mean of the input ± standard deviation. **F)** A cartoon showing the details and coordinates of the genomic region assessed in the ChIP analysis of PEG3 binding to the *Ppox* promoter region.

**Figure S6**

*Snord116* is highly expressed in the hypothalamus, which is one of the brain regions that plays a pivotal role in controlling the sleep-wake cycle. Specifically, in the lateral hypothalamus (LH), there are two groups of neurons, melanin-concentrating hormone (MCH) neurons and orexin (OX) neurons, which exert antagonistic actions on the sleep-wake cycle. MCH neurons promote sleep and maximally fire during REM sleep, while OX neurons promote wakefulness and suppress REM sleep; both contribute to the sleep-wake switch. MCH and OX neurons are important in thermoregulation, which is tightly integrated with the sleep-wake cycle. REM sleep increases when the need for thermoregulatory defence is minimized in TNZ conditions. Our model suggests that Snord116 loss induces a 60% reduction in OX neurons in the LH, while MCH neurons located near OX neurons are unaffected, resulting in an imbalance between MCH and OX neurons. This imbalance between these two types of neurons may explain the sleep alterations and REM sleep disturbances observed in our model, which are also characteristic of PWS syndrome. Thus, the model suggests that *Snord116* regulates REM sleep expression induced by the TNZ via orexin neurons, resulting in an altered thermoregulatory response.
